# Enzymatic Hydroxylation of Aliphatic C-H Bonds by a Mn/Fe Cofactor

**DOI:** 10.1101/2023.03.10.532131

**Authors:** Magan M. Powell, Guodong Rao, R. David Britt, Jonathan Rittle

## Abstract

Manganese cofactors activate strong chemical bonds in many essential enzymes. Yet very few manganese-dependent enzymes are known to functionalize ubiquitous carbon-hydrogen (C-H) bonds, and those that catalyze this important reaction display limited intrinsic reactivity. Herein, we report that the 2-aminoisobutyric acid hydroxylase from *Rhodococcus wratislaviensis* requires manganese to functionalize a C-H bond possessing a bond dissociation enthalpy (BDE) exceeding 100 kcal/mol. Structural and spectroscopic studies of this enzyme reveal a redox-active, heterobimetallic manganese-iron active site that utilizes a manganese ion at the locus for O_2_ activation and substrate coordination. Accordingly, this enzyme represents the first documented Mn-dependent monooxygenase in biology. Related proteins are widespread in microorganisms suggesting that many uncharacterized monooxygenases may utilize manganese-containing cofactors to accomplish diverse biological tasks.

## Main Text

The biochemistry of manganese is intimately linked with dioxygen (O_2_). Nearly all of the O_2_ in the atmosphere was generated via water oxidation at a manganese-containing cofactor in Photosystem II (Fig 1A) (*1*). This enzyme was largely responsible for the Great Oxygenation Event that enabled the emergence of multicellular life (*2*). Aerobic respiration in these organisms inadvertently generates reactive oxygen species, such as superoxide and peroxide, that are often neutralized by Mn-dependent enzymes. Most mitochondria express a Mn-dependent form of superoxide dismutase to safeguard core respiratory enzymes (*3*) and bacteria lacking access to heme cofactors detoxify hydrogen peroxide with Mn-dependent catalases (*4*).

**Fig. 1.**
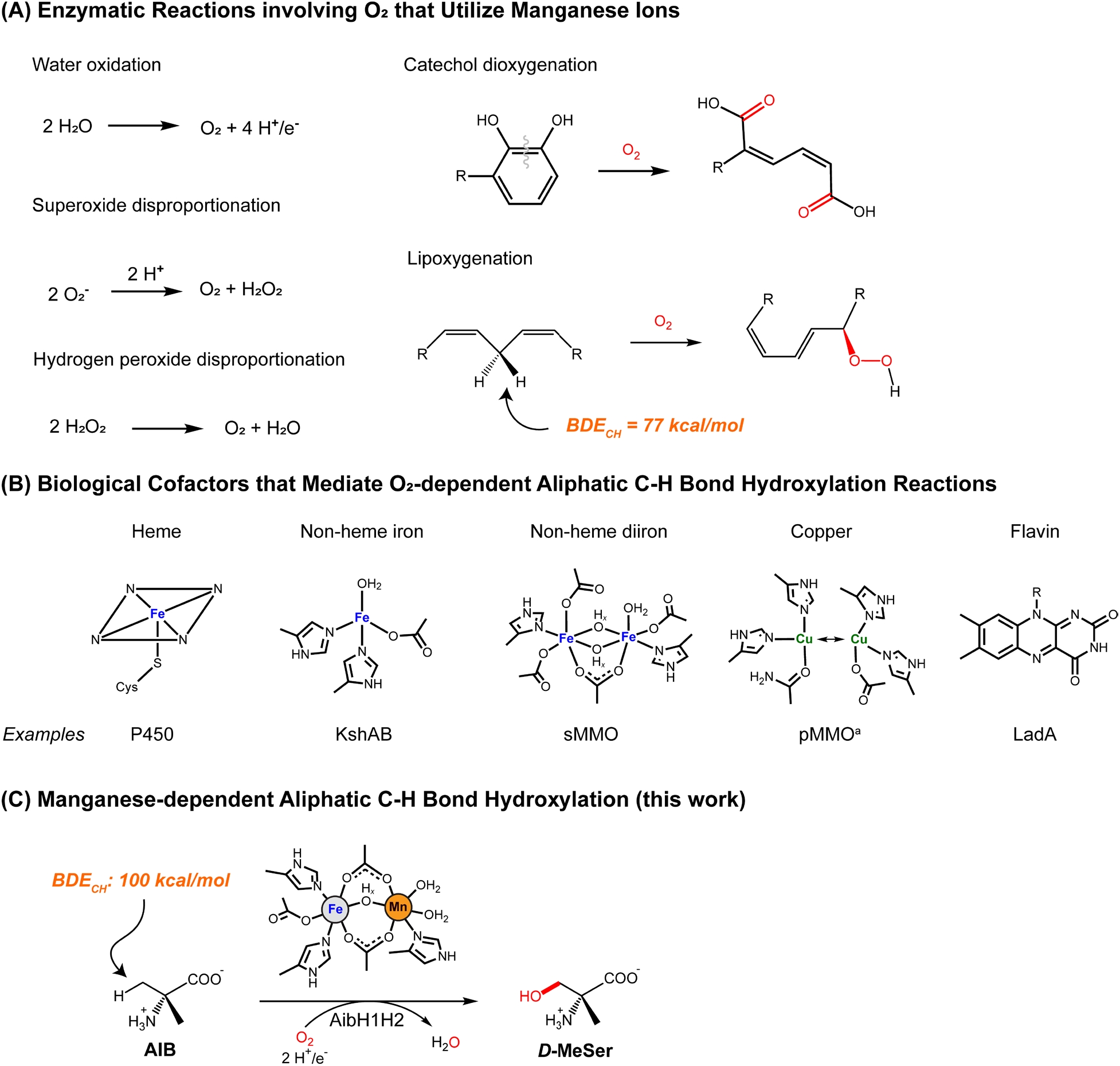
Enzymatic reactions and cofactors relevant to this work. (A) Enzymatic redox reactions involving O_2_ that proceed within a manganese-containing active site. (B) Natural cofactors known to mediate unactivated C*-*H bond hydroxylation. KshAB = 3-ketosteroid-9-alpha-monooxygenase (*17*), sMMO = soluble methane monooxygenase (*18*), pMMO = particulate methane monooxygenase (*19*), LadA = Long-chain alkane monooxygenase (*20*). (C) The enzymatic reaction catalyzed by AibH1H2 (25) and its active site cofactor described in this work. ^a^Copper is the only metal essential for enzymatic activity in pMMO, but the metal nuclearity and precise coordination environment have yet to be reliably determined (*19*).

Given these roles, it is perhaps surprising that very few Mn enzymes are known to use O_2_ to functionalize other substrates. A few ring-cleaving dioxygenases coordinate O_2_ at a Mn site before its insertion into catechol substrates (*5*,*6*) and some fungi employ manganese lipoxygenase to generate reactive lipid metabolites (*7*,*8*). In these latter enzymes, a mononuclear Mn center selectively activates a weak C-H bond (BDE_CH_ = 77 kcal/mol) to generate a radical intermediate that is captured by molecular O_2_. Stronger C-H bonds are not activated. Many bacterial ribonucleotide reductases require a peripheral Mn_2_ or Mn/Fe site to activate O_2_, but these metals do not directly participate in the bond breaking/making reactions occurring at the ribose substrate (*9*–*11*). Besides these enzymes, there are only two other Mn-containing proteins that use O_2_ to mediate C-H bond activation processes in a *stoichiometric* fashion (*12*,*13*).

Of course, many non-Mn enzymes use O_2_ to activate strong, aliphatic C-H bonds. Nature has evolved a large and diverse repertoire of hydroxylases for xenobiotic detoxification (*14*), regulation of cellular function (*15*), and the construction of bioactive natural products (*16*). Yet the embedded cofactors that carry out these chemical processes are exclusive: heme, nonheme iron, flavin, and copper centers are the only biological cofactors known to functionalize aliphatic C-H bonds (Fig 1A) (*14*,*17*–*20*). Accordingly, our foundational understanding of aerobic C-H bond activation stems from the chemical mechanisms operative at this limited set of cofactors.

The absence of natural Mn hydroxylases implies that this metal ion is either not competent to mediate hydroxylation reactions or that such enzymes have eluded discovery. The former explanation is challenged by the numerous *synthetic* catalysts which harness manganese to mediate challenging oxidative chemical reactions including olefin epoxidation, aliphatic C-H bond hydroxylation, and halogenation (*21*–*24*). In this report we demonstrate that the activity of a *Rhodococcus* hydroxylase exhibits a strict dependence on manganese and specifically utilizes an unusual heterometallic Mn/Fe cofactor to effect the catalytic functionalization of an unactivated, primary C-H bond (Fig 1C).

### Establishing the metal dependence on AibH1H2 enzymatic activity

Metabolism of 2-aminoisobutyric acid (AIB) in *Rhodococcus wratislaviensis* proceeds via the initial, selective hydroxylation of the pro-(R) methyl group by the monooxygenase AibH1H2 to furnish α-methyl-*D*-serine (*D*-MeSer) (*25*). Previously, recombinant expression of AibH1H2 in *Escherichia coli* enabled its structural characterization and revealed distinct mono- and di-nuclear metal binding sites housed within the AibH1 and AibH2 protein subunits, respectively (Fig 2A). Metal analyses performed on these samples suggested the presence of iron and zinc ions, leading the authors to propose a structural, mononuclear zinc site in AibH1 and a diiron site in AibH2 that serves as a locus for hydroxylation. Consistent with the latter proposal, many diiron hydroxylases are known (*18*,*26*). The AibH2 subunit is also structurally related to the dinuclear hydroxylase, PtmU3, which displayed superior enzymatic activity in the presence of exogenously supplied iron ions (*27*). But the activity of recombinant AibH1H2 derived from *E. coli* was not determined. Instead, conversion of AIB to *D*-MeSer was only observed via whole-cell recombinant expression of AibH1H2 in *Rhodococcus erythropolis*. These discrepancies in enzyme preparation motivated our independent determination of the active metalated form of AibH1H2.

**Fig. 2.**
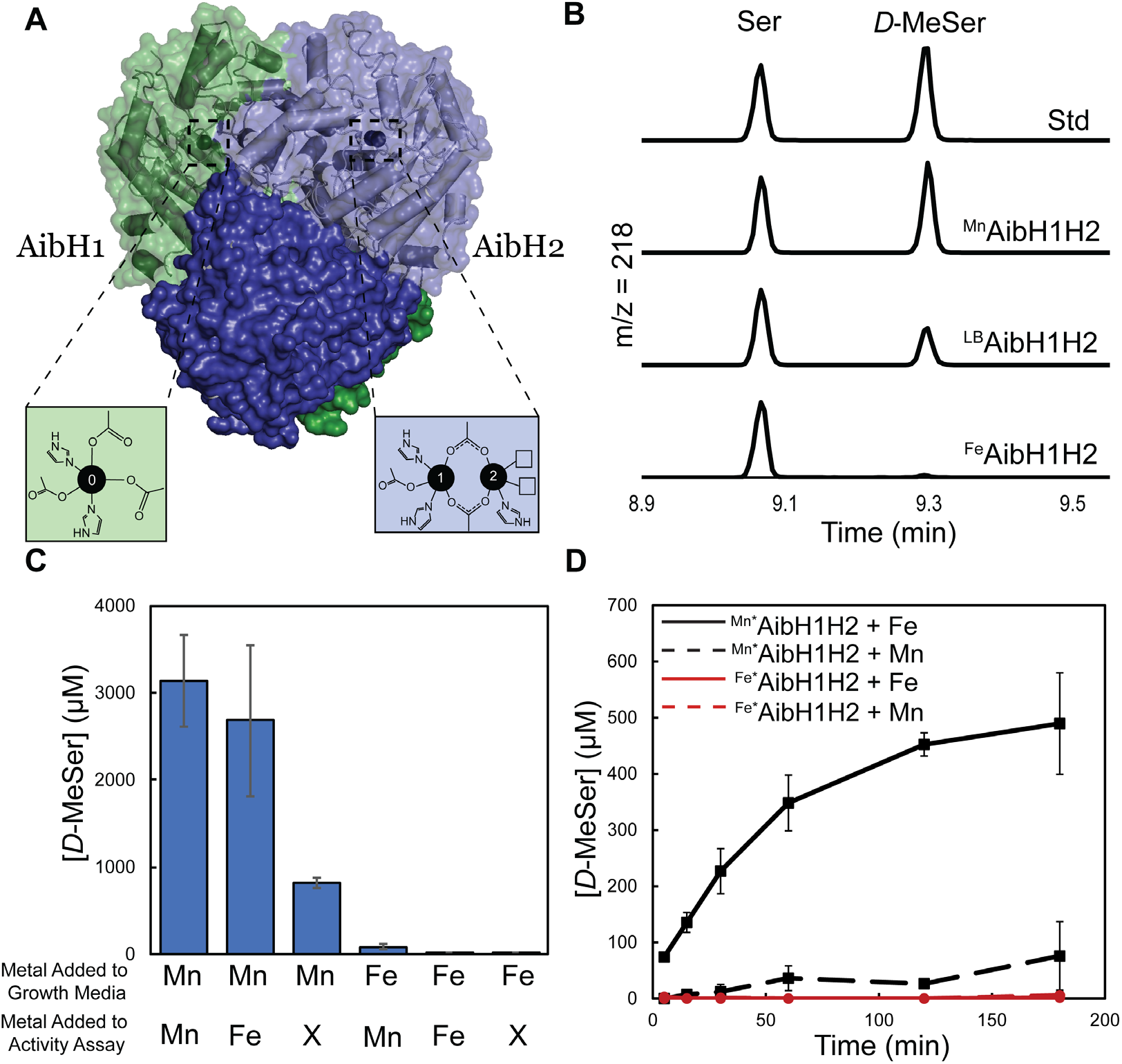
Catalytic activity of AibH1H2 under different metalation conditions. (A) Cartoon and surface representation of the AibH1H2 (αβ)_2_ heterotetramer, made up of AibH1 (green) and AibH2 (blue) subunits alongside their respective metal binding sites. (B) GC-MS chromatograms filtered at m/z = 218 of representative catalytic activity assays of AibH1H2 under different growth conditions with no additional metal added during the activity assay. (C) Catalytic activity of ^Fe^AibH1H2 and ^Mn^AibH1H2 (Table 1, entries 1 and 3, respectively) with one additional equivalent of metal, or an equal volume of water (denoted with X), added to the reaction mixture. (D) Time dependent generation of D-MeSer by ^Mn^*AibH1H2 (black traces, Table 1 entry 4) and ^Fe^*AibH1H2 (red traces, Table 1 entry 5) with one additional equivalent of Mn (dashed trace) or Fe (solid trace) added to the reaction mixture.

We expressed AibH1H2 in *E. coli* grown in M9 minimal medium supplemented with excess Fe^II^, and subsequent purification (Supplemental Methods) furnished protein samples (^Fe^AibH1H2) containing ~3 equiv Fe per AibH1H2 heterodimer and minimal heterometal content (Table 1, entry 1). Fe occupancy in both the mononuclear (Site 0) and dinuclear (Sites 1 and 2) metal sites was evident by inspection of the anomalous dispersion maps obtained from X-ray diffraction experiments performed on single crystals of ^Fe^AibH1H2 (Fig S1). AibH1H2 was found to be enzymatically inactive upon exclusive incorporation of iron ions. Initial assays to determine the enzymatic activity were performed by mixing ^Fe^AibH1H2 with the AIB substrate and sodium ascorbate as a sacrificial reductant under aerobic conditions. Subsequent workup and gas chromatography–mass spectrometry (GC-MS) analyses of these solutions revealed negligible *D*-MeSer production (Fig 2B) (Supplemental Methods). To probe whether the absence of monooxygenase activity was caused by the choice of sacrificial reductant, other commonly utilized reducing systems (*28*,*29*) (e.g., NADH/Phenazine, sodium dithionite, alpha-ketoglutarate) were explored, but none proved competent to produce measurable quantities of *D*-MeSer with ^Fe^AibH1H2 (Fig S2). These results suggested that the active form of AibH1H2 could not be produced under routine expression and purification conditions typically used for nonheme iron enzymes.

**Table 1.** Metal content and reactivity of different AibH1H2 preparations.

| Entry |  | Metal/heterodimer |  |  |  | [D-MeSer]/[AibH1H2]<sup>#</sup> |
| --- | --- | --- | --- | --- | --- | --- |
|  |  | Fe | Mn | Ni | Zn |  |
| 1 | <sup>Fe</sup>AibH1H2 | 2.8 ± 0.1 | 0.08 ± 0.04 | 0.03 ± 0.03 | 0<sup>†</sup> | <2<sup>‡</sup> |
| 2 | <sup>LB</sup>AibH1H2 | 1.67 ± 0.07 | 1.05 ± 0.01 | 0.055 ± 0.003 | 0.131 ± 0.004 | 33 ± 1 |
| 3 | <sup>Mn</sup>AibH1H2 | 0.4 ± 0.1 | 2.2 ± 0.4 | 0.03 ± 0.07 | 0 | 110 ± 30 |
| 4 | <sup>Mn*</sup>AibH1H2 | 0 | 2.4 ± 0.3 | 0 | 0 | 50 ± 10 |
| 5 | <sup>Fe*</sup>AibH1H2 | 1.8 ± 0.2 | 0 | 0 | 0 | <2 |
| 6 | Mn/Fe XRD | 0.0 | 1.9 | 1.3 | 0.0 | 30 |
| 7 | EPR | 0.3 | 1.5 | 0.0 | 0.1 | 73 |

Data for entries 1-5 represent the average of at least three independent trials. Error is the standard deviation. Entries 6-7 represent the specific batch of protein used for the application indicated. <sup>#</sup>Catalytic activity assay performed with the addition of one equivalent of Fe<sup>II</sup> relative to AibH1H2 in the reaction mixture. <sup>†</sup>Values less than the limit of detection of the ICP-OES instrument (0.01 ppm) are reported as zero. <sup>‡</sup>Values less than 2 are below the minimum point on the calibration curve for GC-MS analysis of catalytic activity.

The hydroxylation of AIB is dependent upon manganese incorporation into AibH1H2. The expression of AibH1H2 in rich medium (Lysogeny broth) furnished samples (^LB^AibH1H2) that contained ~1.7 equiv Fe and ~1 equiv Mn per AibH1H2 heterodimer (Table 1, entry 2). The incorporation of tightly-bound manganese ions was evident upon inspection of the electron paramagnetic resonance (EPR) spectra of exhaustively dialyzed samples (Fig S3). Unlike the case for ^Fe^AibH1H2, enzymatic assays employing ^LB^AibH1H2 yielded substantial quantities of *D*-MeSer (33 equiv/AibH1H2). Since the protein structures of these two samples were found to be identical (RMSD = 0.34 Å over all atoms; Fig S1B), our findings suggested an apparent requirement for one or more Mn ions in AibH1H2. This hypothesis was confirmed upon expression of AibH1H2 in M9 minimal medium supplemented with excess Mn^II^ instead of Fe^II^ to afford samples (^Mn^AibH1H2) that displayed 3-fold higher specific hydroxylase activity and likewise a higher Mn content (Table 1, entry 3) relative to ^LB^AibH1H2.

The proposal of a Mn-dependent monooxygenase is controversial because previous reports of Mn-dependent hydroxylation have been discredited due to Fe contamination (*30*–*32*). Similarly in our case, the presence of Fe and other contaminating heterometals in the catalytically active as-isolated preparations of AibH1H2 obscured the identity of the most active cofactor (Fig 2C). We thus subjected these protein samples to mild metal chelation protocols (Supplemental Methods) to standardize the metal content. The resultant samples contained either ~2 equiv Mn (^Mn*^AibH1H2) or ~2 equiv Fe (^Fe^*AibH1H2) and minimal contamination with other metals (Table 1, entries 4-5). With these partially metalated samples in hand, enzymatic assays were performed in the presence of an additional equivalent of Fe^II^ or Mn^II^ in order to vary the identity of the third metal site *in situ*. Hydroxylation of AIB occurred exclusively by ^Mn*^AibH1H2 upon addition of one equivalent of Fe^II^ (Fig 2D). No reaction of the ^Fe^*AibH1H2 protein, regardless of metal addition, yielded product above the limit of detection of the analysis, and similarly low activity was observed following exposure of ^Mn^*AibH1H2 to an additional equivalent of Mn^II^ (Fig 2D). Collectively, these results indicated that the incorporation of two manganese ions and one iron ion afforded the most active form of AibH1H2.

### Characterization of the AibH1H2 cofactors

To determine the locations of these metal ions, crystallographic studies were performed on active preparations (Table 1, entry 6) of AibH1H2. A single crystal grown from a sample of ^Mn*^AibH1H2 premixed with 2 equiv Fe^II^/AibH1H2 was examined by multi-wavelength X-ray crystallography. The unique locations of metal ions were determined by analysis of the anomalous X-ray dispersion collected at either the Mn K-edge or from an anomalous isomorphous difference map stemming from diffraction data collected above and below the Fe K-edge (*33*). Two AibH1H2 heterodimers are found in the asymmetric unit. Both AibH1 subunits display strong anomalous scattering at the Mn K-edge above background levels (Table S1) and lack observable density at the Fe K-edge difference map (Fig S4A) at Site 0, supporting exclusive Mn binding at these sites (Fig 3A). In contrast, the dinuclear sites in AibH2 appear in two distinct metalated forms. View 1 of AibH2 (Fig 3B) contains a dimanganese site with a fully occupied Site 1 metal and partially occupied (~60%) Site 2 metal. View 2 of AibH2 contains two fully occupied sites with predominant Fe occupancy in Site 1 and exclusive manganese occupancy in the solvent-exposed Site 2 (Fig 3C). Since this latter view represents the only location of observable Fe content throughout the structure, we hypothesize that the bimetallic arrangement of View 2 represents the correctly-metalated form of AibH1H2 that participates in substrate hydroxylation.

**Fig. 3.**
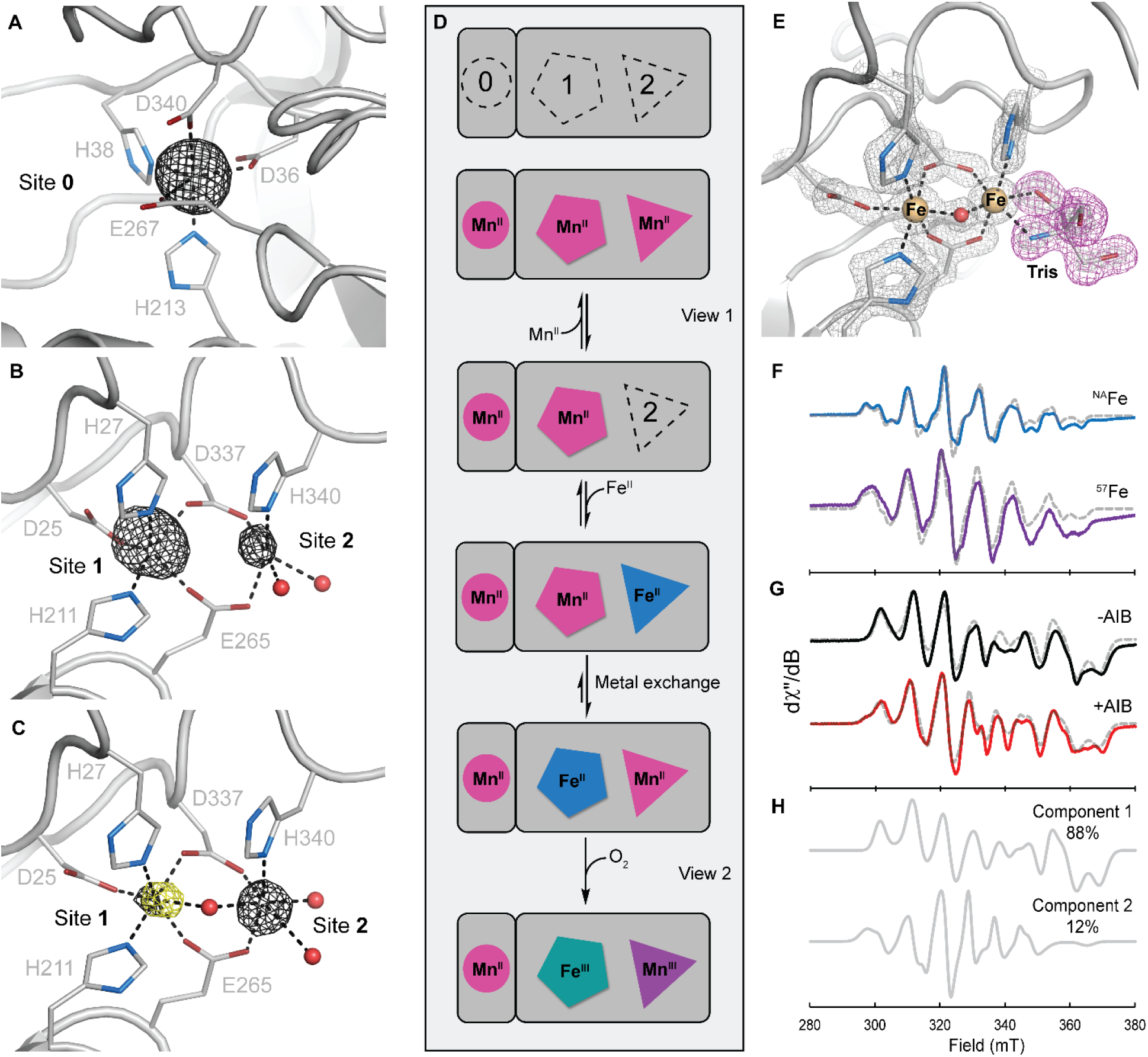
Characterization and proposed metalation of the AibH2 Mn/Fe cofactor. Cartoon representations of the ^Mn^*AibH1H2 protein structure at Site 0 (**A**), View 1 **(B)**, and View 2 **(C)** of the dinuclear cofactor are shown overlayed with the anomalous density difference map at the Mn K-edge (black) and the dual-wavelength isomorphous anomalous density difference map near the Fe K-edge (yellow) each contoured at 5.0σ. (**D**) Proposed metalation scheme of AibH1H2. (**E**) High-resolution crystal structure of ^Fe^AibH1H2 crystallized in the presence of Tris. 2Fo-Fc electron density is contoured at 2.5σ and shown in gray mesh. (**F**) Continuous wave X-band EPR spectra of ^Mn^AibH1H2 in 1 M Tris with one additional equivalent of natural abundance Fe (top, blue) or ^57^Fe (bottom, purple) collected at 15 K and 20 mW power. (**G**) EPR spectra of ^Mn^AibH1H2 in CHES without (top, black) and with (bottom, red) 100 mM AIB taken at 12 K and 63.25 mW power. Simulations are shown in gray dashed lines. (**H**) The extracted components used to simulate the CHES / AIB sample (G, red). Component 1 has identical parameters as the 20 mM CHES samples without AIB (G, black) and Component 2 represents a new species. Refer to the Supplemental Material for simulated spin Hamiltonian parameters. Tris = tris(hydroxymethyl)aminomethane, CHES = N-cyclohexyl-2-aminoethanesulfonic acid.

These crystallographic snapshots allowed us to formulate a hypothesis for cofactor assembly in AibH1H2. In the presence of high Mn^II^ concentrations, this ion will bind to Site 0 and to one or both sites of the AibH2 dinuclear cofactor (Fig 3D). Fe^II^ ions are expected to displace Mn^II^ ions in a mixed Fe^II^/Mn^II^ environment as they form stronger metal-ligand bonds (*34*). However, slow metal dissociation kinetics may prevent observable metal exchange, a possible explanation for the substitutionally inert nature of the mononuclear Mn^II^ site in AibH1 upon exposure to Fe^II^. Fe^II^ is also expected to displace one or both Mn^II^ ions of the dinuclear site, but in the presence of *limiting* Fe^II^ and under thermodynamic control, this ion is expected to bind to the most tightly chelating site. In this scenario, Site 1 is the expected and observed position for Fe^II^ binding, as the coordination environments of Sites 1 and 2 contain five and three amino acid-derived ligands, respectively. This proposed mechanism is similar to that proposed for the Mn/Fe cofactor in class Ic ribonucleotide reductase but distinct from that of R2lox (*9*,*35*–*37*). In cellular contexts, the concentrations of Mn^II^ and Fe^II^ present in the cytoplasm during protein synthesis will influence the outcome of the final metalated state of AibH1H2. These metal ion concentrations can vary widely and are dependent on numerous factors. For example, *E. coli* maintains ~100 μM Fe^II^ and ~10 μM Mn^II^ in its cytoplasm, whereas *Bacillus subtills* is known to possess a 10-fold higher concentration of Mn^II^ under similar growth conditions (*11*). We speculate that *R. wratislaviensis* must possess a relatively high Mn^II^ concentration to allow proper construction of the active heterobimetallic cofactor in AibH1H2. Alternatively, endogenous chaperone proteins (*38*) may selectively deliver manganese ions to the pertinent metal sites during protein synthesis, although experimental data in support of either proposal is not presently available.

Available crystallographic and analytical data strongly support the presence of Mn and Mn/Fe cofactors in the active form of AibH1H2, but the locus of redox activity necessary for O_2_ and substrate activation remains to be determined. Mutations to one or more of the metal binding residues in either AibH1 or AibH2 rendered the protein insoluble or led to a complete loss of enzymatic activity, suggesting that both cofactors play essential roles in the maintenance of structural integrity and/or catalytic activity. We disfavor Site 0 as the site of AIB hydroxylation owing to its sterically occluded nature (Fig S4B). In contrast, the AibH2 dinuclear site is solvent-exposed. A 1.5 Å X-ray crystal structure of ^Fe^AibH1H2 grown in the presence of Tris (Fig 3E) illustrates that exogenous small molecules can access and coordinate to the Site 2 metal.

EPR spectroscopy was used to recapitulate this ligand-bound structure in solution and determine the reactivity of Mn/Fe-metalated AibH1H2 with O_2_ and small molecules. First, a sample of ^Mn^AibH1H2 was combined with one equivalent of Fe^II^ and Tris and subsequently exposed to air for 1 h. Two distinct paramagnetic species were observed in the resultant EPR spectra of these frozen solutions. An isotropic signal characteristic of a mononuclear *S* = 5/2 manganese ion was predominant at high temperatures and ascribed to a Mn^II^ cofactor in Site 0 and/or adventitiously bound Mn^II^ (Fig S5). This signal could be effectively saturated at high powers and low temperatures to reveal an unobscured view of the second EPR-active species. This latter signal (Fig 3F) contained well-resolved features that are noticeably perturbed when isotopically enriched ^57^Fe was incorporated into the sample. These spectra are similar to the *S*TOT = 1/2, antiferromagnetically coupled Mn(III)-Fe(III) forms of *C. trachomatis* ribonucleotide reductase and R2lox (*39*,*40*). Indeed, spectral simulations revealed ***g*-** (2.03 2.03 2.02) and ^55^Mn hyperfine (***A***_Mn_ = [300 250 380] MHz) tensors consistent with the respective spin and oxidation state assignments. The isotropic ^57^Fe hyperfine tensor (***A***_Fe_ = [−70 −70 −70] MHz) further supported the assignment of a high-spin Fe(III) oxidation state. These experiments confirm that the AibH2 cofactor is redox-active and readily oxidized by O_2_ to generate a Mn(III)-Fe(III) redox state.

Since direct substrate coordination to a metal center is observed in other diiron oxygenases (*41*,*42*), we explored the possibility of AIB substrate coordination to the Mn/Fe cofactor in AibH2. The EPR spectrum of AibH1H2 prepared in the absence of coordinating small molecules (Fig. 3G, top) was markedly distinct from that found in the presence of Tris. Most notably, low-field features are lost (<300 mT) and new features emerge at higher fields (~370 mT), and these spectra hence required a distinct set of Hamiltonian parameters for their effective simulation (Table S2). Inclusion of 100 mM AIB to similarly prepared samples resulted in a spectrum displaying subtle, but reproducible perturbations (Fig. 3G, bottom). This EPR signal could only be simulated as a composite of two species with a new 12% component ascribed to an AIB-bound form. The parameters of the this component resembles that of AibH1H2 in the presence of Tris suggesting that both AIB and Tris perturbed the spin center in similar manners (*12*). We propose that AIB is bound either via direct coordination to the Site 2 metal (i.e. Mn) or in a noncovalent manner that substantially alters the geometry and/or protonation state of one or more coordinated ligands. Together, the redox active core and the coordination of AIB at the dinuclear site strongly support the direct involvement of the heterobimetallic Mn/Fe cofactor in substrate hydroxylation by AibH1H2.

### Identification of a widespread family of AibH2-like enzymes

Owing to the paucity of Mn-dependent monooxygenases, we searched for proteins related to AibH2 to determine the biological distribution of enzymes competent to harbor a similar Mn/Fe cofactor. The amino acid sequence of AibH2 is highly divergent from all of the biochemically characterized enzymes of this protein family (PF04909) that more commonly contain monometallic zinc cofactors and catalyze non-redox reactions (Fig 4A) (*43*–*45*). PtmU3, the other established monooxygenase in this protein family, displays 27% sequence identity or 40% similarity to AibH2 (*27*). A multiple-sequence alignment of the UniProt reference proteome sequences displaying greater than 24% identity to AibH2 identified 555 proteins containing all six amino acid residues found to coordinate the Mn/Fe cofactor in AibH2 (Fig 4B). All of these proteins are presently uncharacterized but predicted to possess protein folds and active site structures similar to AibH2 (Fig 4C). The constitution of the genomic neighborhoods (Fig 4D) surrounding the corresponding genes are distinct from those of AibH1H2 and collectively contain few co-occurring proteins that could be used to infer their precise biological function(s). However, a conspicuous 90% co-occurrence of an adjacent small Rieske protein, and 40% co-occurrence of small molecule permeases provides support that these operons are engaged in the catabolism of unknown small molecules. In particular, the tight genomic association between the AibH2-like proteins with the Rieske proteins is reminiscent of catabolic monooxygenases (e.g., Cytochrome P450, soluble methane monooxygenase) which frequently require a dedicated, endogenous reductase (*14*,*18*). Accordingly, we postulate that these 555 uncharacterized PF04909 proteins represent members of a new class of monooxygenases that harbor Mn/Fe cofactors to catalyze as yet unknown hydroxylation reactions. It is noteworthy that a large portion of these uncharacterized proteins stem from organisms known to display high cytoplasmic manganese concentrations, including members of *Bacilli* (*11*), radiation and/or desiccation resistant bacteria (*46*,*47*), and halophilic microorganisms (*48*). Experimental efforts to establish the enzymatic reactivity and cofactor content of these candidate monooxygenases are ongoing in our laboratories.

**Fig. 4.**
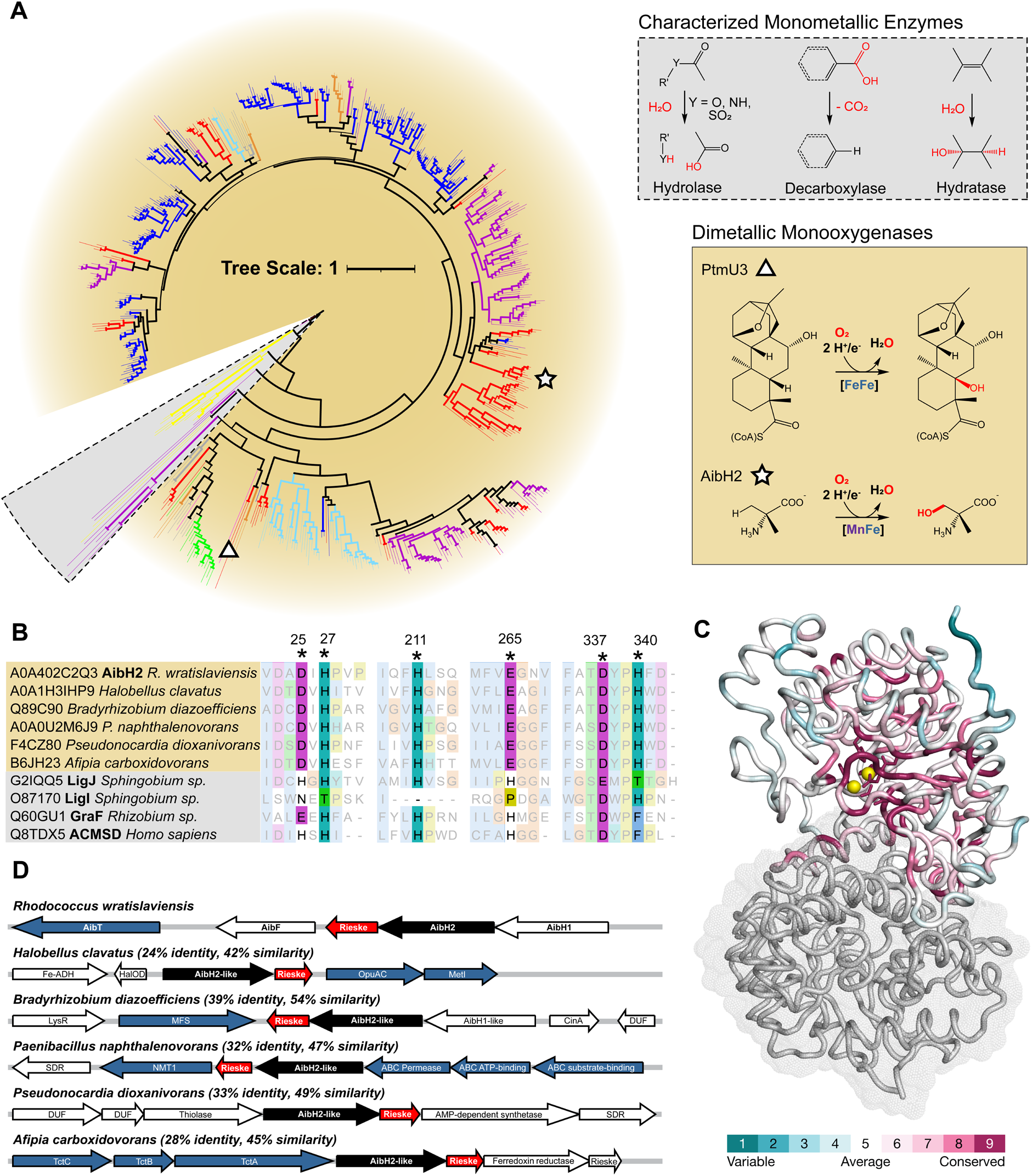
Bioinformatic identification of an uncharacterized family of AibH2-like proteins. (**A**) Unrooted, neighbor-joining phylogenetic tree of 555 Uniprot Reference Proteome sequences with at least 24% identity to AibH2 and conserved metal-coordinating residues. Biochemically-characterized representatives of the PF04909 protein family were included for context. Branch thickness is proportional to bootstrap values. The branches are colored according to the taxonomic group: Red: Actinobacteria; Magenta: Proteobacteria; Blue: Bacilli; Orange: Chloroflexi; Yellow: Eukaryote; Teal: Haloarchaea; Green: Cyanobacteria; Black: Other. The underlying shading reflects the enzymatic reactions (inset) known or expected to be catalyzed by these enzymes. (**B**) Multiple sequence alignment of representative candidate monooxygenase sequences and characterized monometallic enzymes highlighting the metal binding residues found in AibH2. (**C**) Ribbon and surface representation of AibH1H2 illustrating the localization of variable (cyan) and conserved (purple) regions of 300 randomly chosen AibH2-like sequences. (**D**) Genome neighborhood diagrams of AibH2 and AibH2-like proteins. Genes are colored according to inferred function: Black: AibH2-like monooxygenase; Red: Rieske-type ferredoxin; Blue; small molecule permease components.

## Conclusions

The *in vitro* studies of AibH1H2 presented in this work provide the first unambiguous evidence for Mn-dependent hydroxylation in Nature. The available reactivity, crystallographic, and spectroscopic data collectively support a redox-active Mn/Fe cofactor that can activate O_2_ and bind AIB *en route* to its hydroxylation. The presence of manganese at the key site of AIB coordination (Site 2) emphasizes its critical role in the functionalization of a strong aliphatic C-H bond. Accordingly, this unusual Mn/Fe cofactor exhibits reactivity on par with the highly reactive Fe- or Cu-dependent hydroxylase active sites (Fig 1B). Our results expand the known roles of manganese in biology and motivate further studies to understand how and where Mn-dependent monooxygenases function.

## Acknowledgments

We thank members of the Rittle group, M. Green, D. Newman, J. Peters and F. A. Tezcan for discussions. We also thank the Lawrence Berkeley National Laboratory Catalysis Center at UC Berkeley for the use of GC-MS instrumentation. Parts of this research were carried out at the Stanford Synchrotron Radiation Lightsource (supported by the DOE, Office of Basic Energy Sciences contract DE-AC02-76SF00515 and NIH P30-GM133894) and the Advanced Light Source (supported by the DOE, Office of Basic Energy Sciences contract DE-AC02-05CH11231and NIH P30-GM124169-01).

## Author contributions

Conceptualization: MMP, JR
Methodology: MMP, GR, JR
Investigation: MMP, GR, RDB, JR
Visualization: MMP, JR
Supervision: RDB, JR
Writing: MMP, GR, RDB, JR

## Data and materials availability

All structures were validated and deposited in the Protein Data Bank (PDB) under the following accession numbers: 8FUL, AibH1H2 expressed from Lysogeny Broth; 8FUM, Fe-metalated AibH1H2 with bound Tris; 8FUN, Mn/Fe metalated AibH1H2; 8FUO, Fe-metalated AibH1H2. All other data are available in the manuscript or the supplementary materials.

## Supplementary Materials

Materials and Methods

Figs. S1 to S10

Tables S1 to S7

References (*49*–*61*)

