## Supplemental Materials for "Enzymatic Hydroxylation of Aliphatic C-H Bonds by a Mn/Fe Cofactor"

**The PDF file includes:**

Materials and Methods
Figs. S1 to S10
Tables S1 to S7
References (*49-61*)

### Materials and Methods

General

All chemicals were purchased from commercial suppliers and used directly as received unless noted otherwise. The plasmid pACYC-GroEL/ES-TF was a gift from Karl Griswold (Addgene plasmid #83923; http://n2t.net/addgene:83923; RRID:Addgene 83923). Metal analysis was conducted in the Microanalytical Facility in the College of Chemistry at the University of California, Berkeley using a Perkin Elmer ICP Optima 7000 DV Spectrometer. All degassed chemicals were rendered anaerobic by equilibration overnight in a Coy anaerobic chamber equipped with a 1.5-2.5% $H_2/N_2$ environment and a palladium catalyst. All aqueous solutions were prepared using water purified by a MilliQ Academic water purifier and exhibited a conductivity of 18.2 MΩ. Genetic sequencing was performed at the University of California, Berkeley DNA Sequencing Facility. Large-scale (>0.5 L) bacterial cultures were grown using a LEX-48 bioreactor (Epiphyte3). An Agilent 7890 gas chromatography-mass spectrometry (GC-MS) system equipped with an HP-5MS Agilent column (30m x 0.250 mm x 0.25 μm) at the Lawrence Berkeley National Laboratory Catalysis Laboratory was used for GC-MS analysis. Nuclear Magnetic Resonance (NMR) spectra were collected on a Bruker AVQ-400 spectrometer at the University of California, Berkeley College of Chemistry NMR Facility.

Construction of Vectors and Strains for Over-Expression of AibH1H2 from *Rhodococcus wratislaviensis* in *Escherichia coli*

Multiple expression vectors for AibH1H2 were generated to enable straightforward protein synthesis or to ameliorate irreproducible protein yields and/or the ability to obtain high quality single crystals. The specific utility and construction for each individual vector is described below and summarized in Table S5.

The $His_6$-affinity-tagged pACYC-AibH1H2 plasmid was used to obtain single crystals of $^{Fe}$AibH1H2 and was constructed as follows. Codon-optimized genes of AibH1 (UniProt KB accession no. A0A402C2V4) and AibH2 (UniProt KB accession no. A0A402C2Q3) were ordered from Twist Bioscience (Table S3). The AibH1 gene is preceded by the NcoI restriction enzyme cut site, a $His_6$-tag, and the TEV protease recognition sequence, and it is followed by the HindIII restriction enzyme site on the 3' end. The AibH2 gene is flanked by the NdeI (5') and XhoI (3') restriction enzyme recognition sequences. Synthesized gene fragments were resuspended in 50 μL of nuclease-free Millipore water and frozen at -20 °C prior to usage. The pACYC-GroEL/ES-TF vector (Addgene plasmid #83923) and AibH1 gene fragment were each digested with NcoI and HindIII. The gene fragment and vector were purified using a PCR and gel cleanup kit (Biomiga), respectively. The 1.2 kilobase pair (kb) AibH1 fragment was ligated into the digested vector (5.2 kb) with T4 ligase and buffer following the New England Biolabs (NEB) protocol and transformed into NEB Turbo chemically competent *E. coli* for amplification. The plasmid was extracted using a plasmid extraction kit (Biomiga). The resulting plasmid and the 1.2 kb AibH2 gene fragment were digested with NdeI and XhoI, ligated, and transformed into NEB Turbo *E. coli*, as described above. Extraction of the resulting plasmid yielded the pACYC-AibH1H2 plasmid with chloramphenicol resistance, and the complete sequence of both genes was confirmed by plasmid sequencing using the primers LacI-F, pAC-Md-F, T7 term, and pAC-T7-1R (Table S4). pACYC-AibH1H2 was then transformed into BL21(DE3) competent *E. coli* for protein overexpression.

To increase protein expression yield, the AibH1 and AibH2 genes were moved to the pET-Duet vector with ampicillin resistance, allowing the genes to be co-expressed with the pACYC-

GroEL/ES-TF plasmid (Addgene plasmid #83923), as the chaperone proteins GroEL/ES and TF promote efficient protein folding during heterologous expression. The pET-Duet-AibH1H2 plasmid was constructed following the same protocol as described above for the pACYC-AibH1H2 plasmid construction, except starting with the pET-Duet-1 vector (EMD Millipore Corp.). The plasmid was confirmed by sequencing using the following primers: T7 term, T7, DuetDOWN, and DuetUP (Table S4). In the final step, the pET-Duet-AibH1H2 vector was transformed to *E. coli* BL21(DE3) cells pre-transformed with pACYC-GroEL/ES-TF and made chemically competent following a previously reported protocol (*49*). This strain was used for all experiments unless otherwise specified.

A truncated pET-Duet-AibH1H2-Δ17-24 was generated by deletion of base pairs 49-72 of AibH1 from the pET-Duet-AibH1H2 plasmid using the primers AibH1_Δ17-24-F and AibH1_Δ17-24-R (Table S4) in a polymerase chain reaction (PCR). The NEB manufacturer PCR protocol for Phusion polymerase was followed with an annealing temperature of 75 °C. The resulting DNA was digested for one hour with DpnI at 37 °C and 5 μL of the PCR mixture was transformed directly into NEB Turbo *E. coli*. The plasmid was extracted, confirmed by DNA sequencing, and transformed to chemically competent *E. coli* BL21(DE3) cells pre-transformed with pACYC-GroEL/ES-TF as described for pET-Duet-AibH1H2. This strain was used to generate the Mn/Fe-containing crystal structure (8FUN).

Heterologous Protein Expression and Purification

All strains for protein over-expression were cultivated in either commercially available Lysogeny broth (LB, Miller) or M9 minimal medium, as indicated in the main text. M9 medium used herein contains MilliQ water supplemented with 70 mM phosphate ($KH_2PO_4$ / $Na_2HPO_4$), 8.6 mM NaCl, and 18.7 mM $NH_4Cl$. The pH of these solutions was adjusted with NaOH to pH 7.4 and autoclaved followed by the addition of 900 μM $MgSO_4$, 175 μM $CaCl_2$, and 0.375% glucose from filtered stock solutions. The medium was supplemented with 100 mg/L filtered ampicillin for strains containing pET-Duet-AibH1H2 and/or 34 mg/L chloramphenicol (dissolved in ethanol) for strains containing pACYC-AibH1H2 or pACYC-GroEL/ES-TF plasmid. Cultures were grown immersed in a water bath temperature set to 38 °C and vigorously aerated with HEPA-filtered compressed air. At an $OD_{600} \approx 0.6$, the water bath temperature was decreased to 18 °C, and 0.125 mM of divalent manganese or iron, in the form of $MnCl_2 \cdot 4H_2O$ or $(NH_4)_2Fe(SO_4)_2(H_2O)_6$, respectively, and 1 mM isopropyl β-D-1-thiogalactopyranoside (IPTG) were added immediately. The cultures were allowed to grow for 18-24 h before the cells were harvested by centrifugation at 5000 rpm for 5 minutes at 4 °C. The cell pellets were frozen at -80 °C until purification. All purification steps were performed at 4 °C. Thawed pellets from 12 L of growth media were resuspended in lysis buffer (200 mM NaCl, 50 mM HEPES pH 8.0) to a final volume of ~60 mL and sonicated for 10 minutes of total pulse time (3 seconds pulse, 6 seconds off). For larger protein batches, sonication was successively performed in 60 mL fractions and then pooled following cell lysis. The lysate was centrifuged for 1 h at 12,000 rpm to remove cell debris and the resulting supernatant was loaded onto a column equipped with Ni-NTA resin (Thermo Fisher Scientific). The resin was subsequently washed with 100 mL of lysis buffer, washed further with 600-750 mL of wash buffer (lysis buffer + 25 mM imidazole), and eluted with 150 mL of elution buffer (lysis buffer + 250 mM imidazole). The protein was concentrated to <50 mL using an Amicon® stirred cell (Millipore Sigma) equipped with a 30 kDa filter and subsequently buffer exchanged into 20 mM HEPES pH 8.0 via two rounds of dialysis against 4 L of buffer (one round was overnight and one for a minimum of 2 h). Protein purity was evaluated via sodium dodecyl sulfate–

polyacrylamide gel electrophoresis (SDS-PAGE, Fig. S9), and the metal content determined using ICP-OES. Protein was stored at 4 °C until further experiments, except where further manipulation was required, described below.

When the pET-Duet-AibH1H2-Δ17-24 plasmid was used, a final step to remove the $His_6$-tag via TEV protease cleavage was required. The purified AibH1H2 protein was stirred in lysis buffer at a final concentration of 1 mg/ml with 2.5 mM dithiothreitol and 1 mg of TEV protease for every 10 mg of AibH1H2 overnight at 4 °C. The solution was passed down a Ni-NTA affinity column, and the flow-through collected and buffer exchanged to 20 mM HEPES pH 8.0 via dialysis as described above.

<u>Metal Chelation and Reconstitution</u>

Protein that required further metal chelation and reconstitution was prepared the same way as described above, except the protein was dialyzed against 4 L of 20 mM HEPES pH 8.0 + 10 mM ethylenediaminetetraacetic acid (EDTA) for 2 h before undergoing the dialysis to 20 mM HEPES pH 8.0. Subsequently, the protein was degassed by equilibration in a Coy anaerobic chamber for at least 1 h. The sample was reduced by the addition of 6 mM sodium dithionite and 20 μM methyl-viologen and allowed to incubate at room temperature for 30 minutes. For Fe-grown protein, contaminating metals were chelated by treatment with 10 equivalents (relative to the concentration of AibH1H2) of EDTA by addition of a degassed 200 mM EDTA stock and allowed to incubate for 1 h at room temperature. For Mn-grown protein, iron was selectively chelated by the addition of 10 equivalents (relative to the concentration of AibH1H2) of ferrozine by the addition of degassed 100 mM ferrozine stock and incubation for 30 minutes at room temperature. *Exposure of protein samples to chelating molecules over longer periods of time results in enzyme preparations resistant to remetallation.* These partially chelated protein samples were then buffer exchanged in the Coy chamber by concentrating the protein to ~5 mL using an Amicon® stirred cell and filling with degassed 20 mM HEPES pH 8.0 up to 100 mL three times. The protein was then reconstituted by addition of five equivalents of $MnCl_2{\cdot}4H_2O$ or $(NH_4)_2Fe(SO_4)_2(H_2O)_6$ to the Mn- or Fe-protein, respectively, and incubated anaerobically overnight at 4 °C. Excess metal was removed by repeating the buffer exchange in the Amicon® stirred cell as described above. Metal content was determined by ICP-OES, and protein was stored at 4 °C until further experiments. The average yield of these combined metal chelation and reconstitution protocols is ~70% (Fe-protein) and ~60% (Mn-protein).

<u>Determination of Protein Concentration</u>

Protein concentration was estimated using the absorbance at 280 nm of protein samples in 20 mM HEPES (NanoDrop 2000c, Thermo Fisher Scientific). The extinction coefficients for AibH1 and AibH2 were predicted with the biochemical analysis tool on the Benchling website and the expected extinction coefficient of AibH1H2 was calculated by taking the average of the constituent monomers (Table S5).

<u>Catalytic Activity Assays and Gas Chromatography-Mass Spectrometry (GC-MS) Analysis</u>

Protein was degassed in a Coy chamber for at least 1 h prior to all enzymatic assays. The final concentration of protein was 25 μM in 100 μL reactions. One equivalent of the appropriate metal from a degassed 2.5 mM metal stock was added to the protein (20 mM HEPES, pH 8.0) and the resultant solution was incubated for 5 min anaerobically, followed by 5 min in air. The reaction was initiated by combining a pre-mixed aerobic reaction mixture of 5 mM ascorbate (100 mM stock), 100 mM Aib (1 M stock), and 20 mM HEPES pH 8.0 (1 M stock) to the protein solution in a 1.5-mL Eppendorf tube. Time-dependent reactions (Fig. 2D) were performed as described except with 200 mM AIB and 1 mM Ser as the internal standard added before initiating the reaction. The reaction tubes were incubated at 30 °C with the cap open and shaken at 300 rpm for 3 h (unless otherwise noted). $^{1}$H NMR spectroscopy (Fig S7) of the resulting filtered solutions can be used to directly observe *D*-MeSer production. For quantitative analyses, the reaction products were silylated and analyzed by GC-MS. Reactions were quenched by the addition of 10 µL 0.4 M trichloroacetic acid, followed by the addition of 2 µL of 50 mM serine to serve as an internal standard for GC-MS analysis. The protein precipitant was pelleted by centrifugation at 13,200 rpm for 5 minutes. 90 µL of the supernatant was transferred to a 600-µL Eppendorf tube, frozen in liquid nitrogen and lyophilized on a Schlenk line. The residue was combined with a 1:1 mixture of *N*-methyl-*N*-(trimethylsilyl)trifluoroacetamide and acetonitrile (50 µL each) and incubated at 70 °C for 30 minutes. The debris was pelleted by centrifugation for 5 minutes and 1 µL of the supernatant was analyzed on by GC-MS (Figs S10-11). The temperature was held at 80 °C for 2 min, and then ramped up to 300 °C at 15 °C/min, and finally held at 300 °C for 10 min. *D*-MeSer production was quantified using the extracted ion chromatogram (EIC) at m/z = 218 divided by the EIC at m/z = 218 of Ser, which served as an internal standard. The yields were determined by comparison to a calibration curve of 100 µL of *D*-MeSer standard subjected to the same experimental conditions and workup as the activity assays. A new calibration curve was prepared at the time of each new experiment.

X-Ray Crystallography

Crystals of all proteins were obtained by sitting-drop vapor diffusion at room temperature in an anaerobic Coy chamber. All solutions and crystal trays (Hampton) were degassed overnight prior to use. The protein was degassed in the Coy chamber for at least 1 h and subsequently incubated with any additional metals for at least 15 minutes prior crystallization. Each reservoir was filled with 200 µL of precipitant solution, and 2 µL of the solution was added to the well containing 2 µL of 6 mg/ml protein. Experimental crystal growth conditions are summarized in Table S6. Good quality crystals generally appeared after 2-4 weeks of maturation. The crystal for the Tris-bound $^{Fe}$AibH1H2 structure was transferred to a well containing 2 µL of the precipitant solution plus 25% PEG400 with 200 µL of this solution in the reservoir and incubated for 14 hours for cryoprotection prior to harvesting and freezing in liquid $N_2$. All other crystals were harvested and frozen in liquid $N_2$ directly. X-ray crystallographic data were collected at the Advanced Light Source (ALS) or the Stanford Synchrotron Radiation Lightsource (SSRL), with the specific beamlines and the energy of data collection tabulated in Table S6. Diffraction data were processed using iMosfilm followed by scaling and merging using Aimless (*50, 51*). Phasing was performed using Phaser-MR via molecular replacement with PDB ID: 6M2I as the initial search model. The final models were generated by refinement in phenix.refine and manual modeling with COOT. Electron density maps were generated in Phenix, and PyMol was used for the generation of all figures. Anomalous difference maps were created with Phenix and iron-specific anomalous DANO maps were created using the ScaleAndMerge and Fobs_minus_Fobs routines within Phenix (*52*). No map sharpening algorithms were utilized in the analysis of anomalous dispersion data.

Electron Paramagnetic Resonance Spectroscopy

EPR studies were performed in the CalEPR center in the University of California, Davis. X-band continuous-wave EPR spectra were recorded on a Bruker Biospin EleXsys E500 spectrometer equipped with a super high Q resonator (ER4122SHQE) in the perpendicular mode. Cryogenic temperatures were achieved and controlled by using an ESR900 liquid helium cryostat with a temperature controller (Oxford Instrument ITC503). Spectrometer settings were: conversion time, 60 ms; modulation amplitude, 0.8 mT; modulation frequency, 100 kHz. The samples in 1 M Tris were collected at 15 K and 20 mW power, and the CHES samples were collected at 12 K and 63.25 mW power. EPR spectra were simulated in Matlab with Easyspin 5.2.28 toolbox (*53*).

The naturally abundant Fe ($^{NA}$Fe) EPR samples were prepared as follows. Samples of as-isolated $^{Mn}$AibH1H2 (in 20 mM HEPES pH 8.0) were degassed by equilibration in a Coy chamber for at least one hour. One equivalent of $(NH_4)_2Fe(SO_4)_2(H_2O)_6$ was added to the protein and incubated at room temperature for 5 min prior to exposure to air. The samples were then buffer exchanged into the appropriate air-saturated buffer (1 M Tris or 20 mM CHES pH 8.9) via five rounds of concentration and dilution in 0.5 mL Amicon® centrifugal unit with a 30 kDa membrane (Millipore). The sample with 100 mM AIB was prepared as described above except the buffer contained 20 mM CHES and 1 M AIB (pH 8.9), and the sample was subsequently diluted 10x with 20 mM CHES pH 8.9 to yield a final concentration of 100 mM AIB in 20 mM CHES pH 8.9. After incubation in air for 1 hour, the samples were frozen in liquid $N_2$ for EPR analysis. Final protein concentrations were 250 µM AibH1H2 (in CHES) or 271 µM AibH1H2 (in Tris).

For the $^{57}$Fe sample in 1 M Tris, a 50 mM $^{57}FeCl_2$ stock was prepared anaerobically in 100 mM HCl and frozen at -80 °C until use. Upon thawing in the anaerobic Coy chamber, the $^{57}FeCl_2$ was diluted 10x to 5 mM using 100 mM HEPES pH 8.0. One equivalent of this 5 mM $^{57}FeCl_2$ stock was added to mildly chelated $^{Mn*}$AibH1H2 (see metal chelation protocol) to minimize contamination of $^{NA}$Fe. The sample was oxidized and buffer exchanged, as described above, prior to freezing in liquid $N_2$.

<u>Bioinformatic Analyses</u>

A BLAST search of the AibH2 amino acid sequence (Uniprot: A0A402C2Q3) on the UniProt Reference Proteome database was used to identify all homologs displaying >25% sequence identity. The resultant sequences were aligned using MUSCLE (*54*) within the UGene software package (*55*). Sequences lacking one or more of the aligned amino acid residues found to coordinate the Mn/Fe cofactor (25D, 27H, 211H, 265E, 337D and 340H in AibH2) were removed, leaving 555 unique proteins. The genome neighborhoods (Fig 3B) of these proteins were downloaded and evaluated using the Enzyme Function Initiative suite of online tools and visualized using Gene Graphics (*56, 57*). To correlate the amino acid conservation with predicted 3D structures, 255 sequences were randomly eliminated, and the resultant 300 sequences were used to generate Fig 3D using the default algorithm parameters on the ConSurf website (*58*).

All resultant 555 amino acid sequences (and AibH2) are classified to the protein family (Pfam) PF04909. To evaluate their evolutionary relatedness to other biochemically characterized enzymes within this family, these sequences were aligned alongside the 30 SwissProt-annotated PF04909 sequences and the sequence for PtmU3. The resultant multi-sequence alignment was manually curated via the elimination of all aligned amino acid positions bearing >50% gaps. This alignment was analyzed using the NGPhylogeny suite of tools (*59*) to create the neighbor-joining, unrooted phylogenetic tree in Fig 4A. PhyML was used to determine the best evolutionary model (LG substitution, empirical equilibrium frequencies, Gamma shape estimated at 1.513) and the tree was constructed with the AIC statistical criterion and NNI tree topology. Branch support values employed the aBayes approximation (*60*) and revealed a high level of confidence across the majority of the tree. The Interactive Tree of Life was used for graphic generation (*61*).

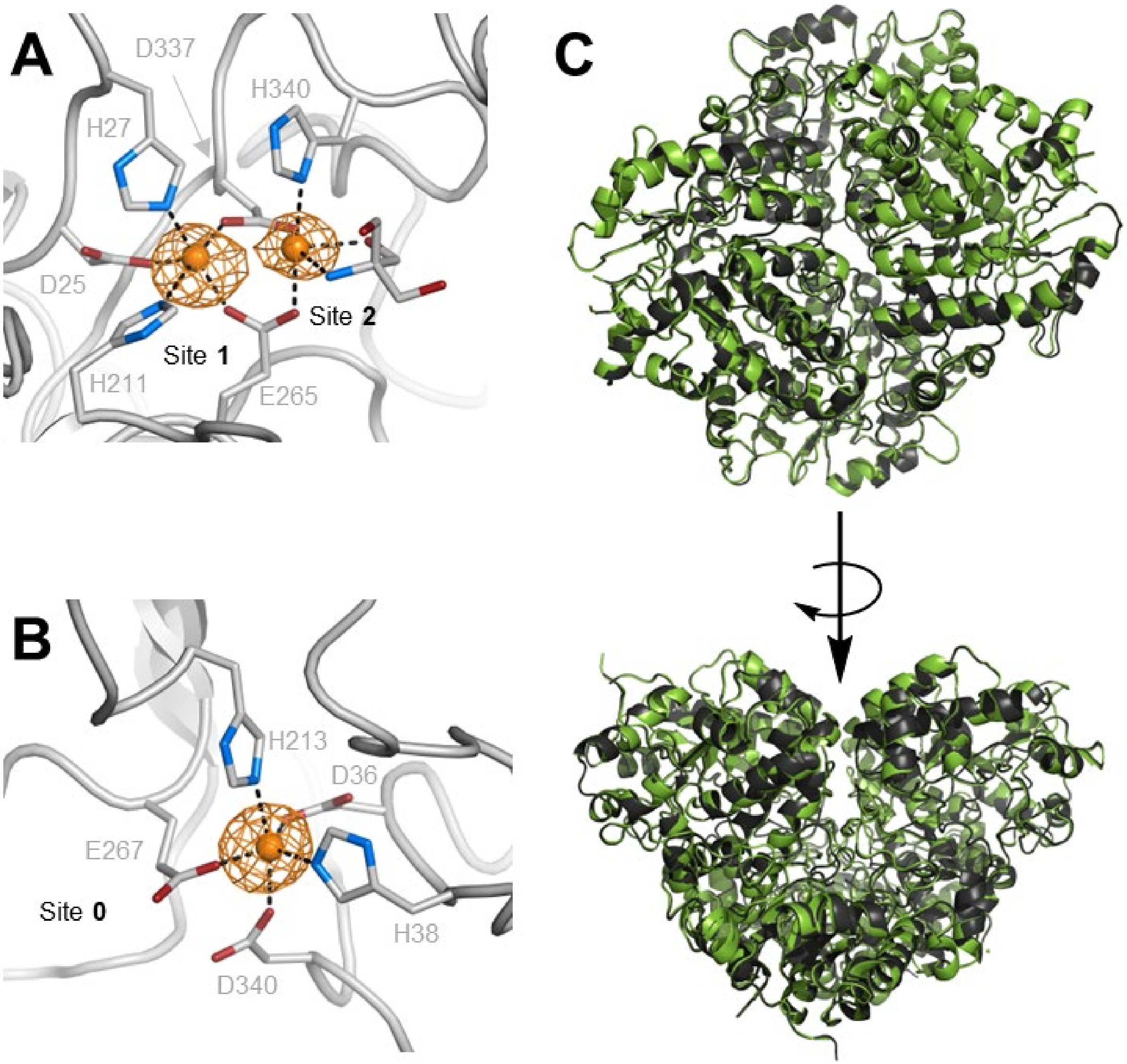


**Figure S1.** Anomalous dispersion density maps (contoured at 11σ) at the (A) dinuclear AibH2 site and the (B) mononuclear AibH1 site. X-ray diffraction data was collected at the Fe K-edge (7132 eV) on a crystal of $^{Fe}$AibH1H2 (8FUO). (C) Structural overlay of $^{LB}$AibH1H2 (green, 8FUL) and $^{Fe}$AibH1H2 (black) (RMSD = 0.34 Å)

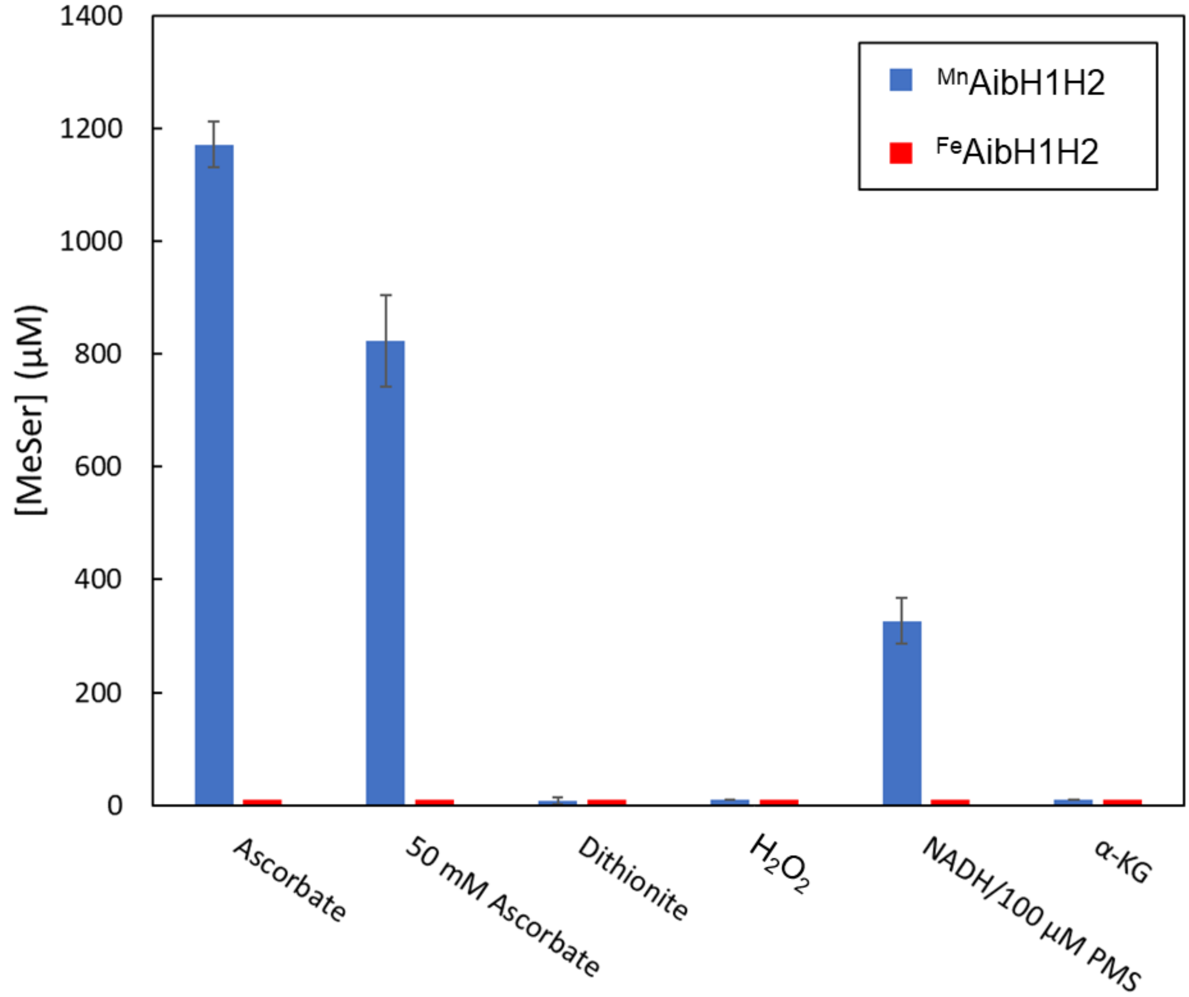


**Figure S2.** *D*-MeSer production from enzymatic assays performed either in the presence of 25 µM $^{Fe}$AibH1H2 (red) or 25 µM $^{Mn}$AibH1H2 (blue), an additional equivalent of $Fe^{II}$, and the listed small molecules to serve as reducing equivalents. Final concentrations of all small molecules were 5 mM unless otherwise indicated. PMS = phenazine methosulfate, α-KG = alpha-ketoglutarate. Each bar is the mean of three independent trials, and the error bars represent the standard deviation in observed *D*-MeSer yield. In all experiments with $^{Fe}$AibH1H2, no *D*-MeSer was observed above the detection limit (~25 µM).

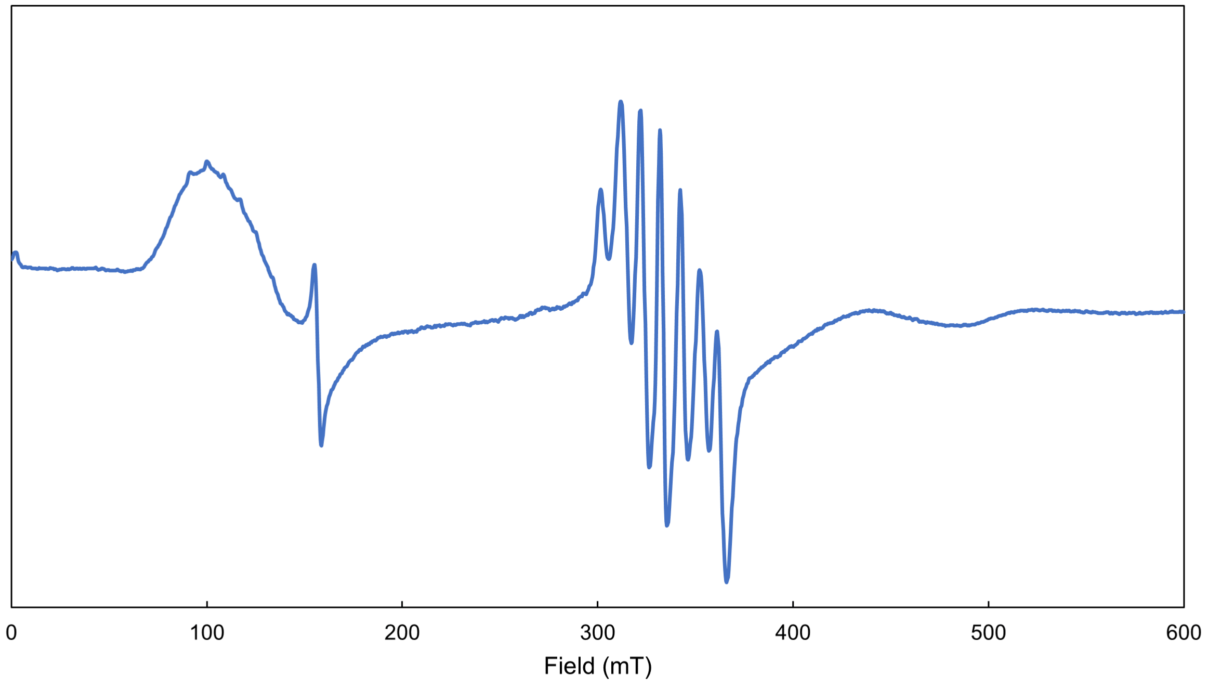


**Figure S3.** X-band CW EPR spectrum of 270 μM $^{LB}$AibH1H2 in 20 mM HEPES buffer, pH 7.5. Spectrum collected at 5 K and 2 mW power.

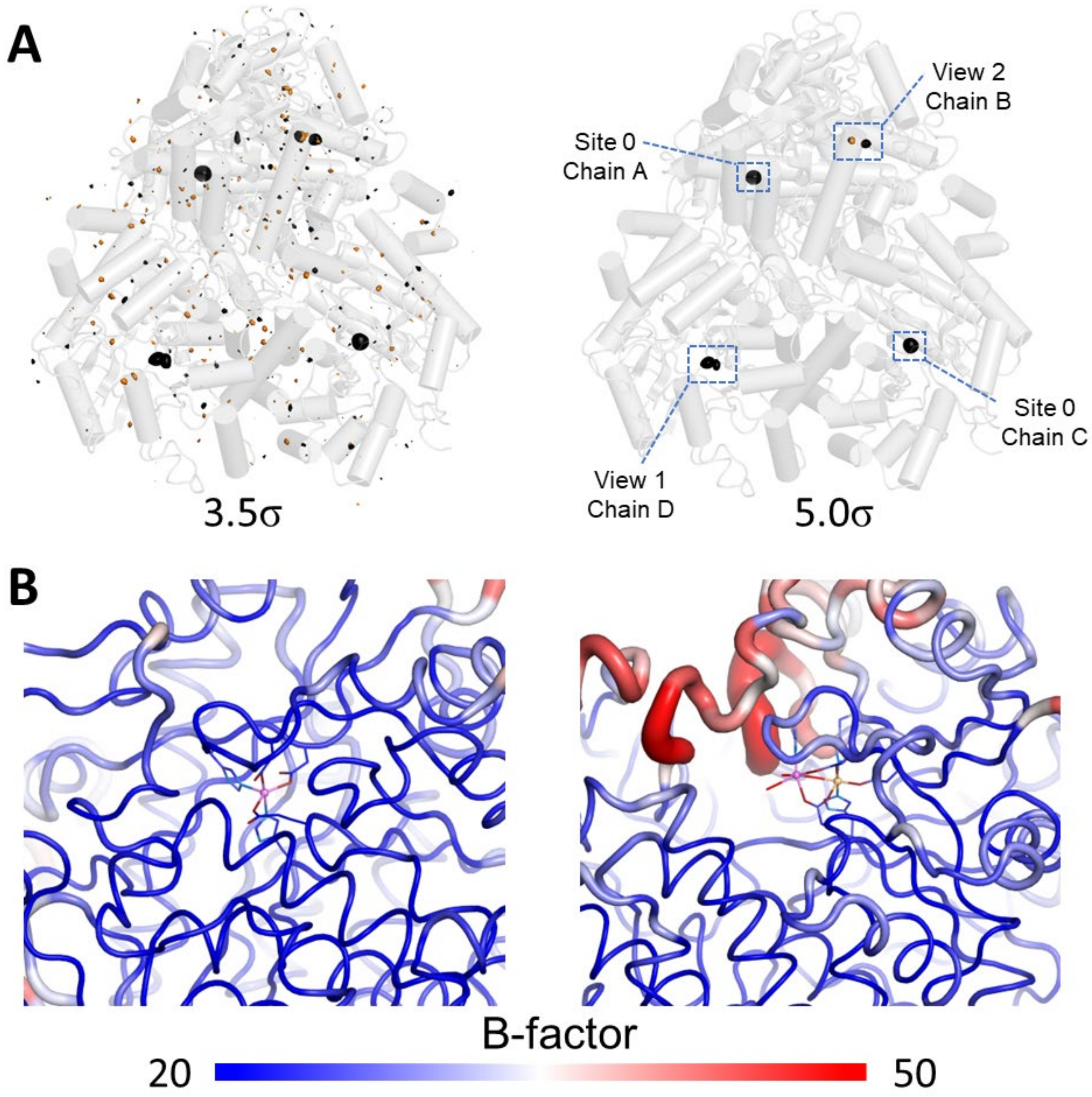


**Figure S4.** (A) Anomalous difference density map at the Mn K-edge (6550 eV, black surface) and Fe-specific dual-wavelength anomalous density difference map (7092 and 7132 eV) each drawn at 3.5σ (left) and 5.0σ (right) for structure 8FUN. The labels correspond to the specific metal binding sites discussed in the main text. (B) Cartoon representations of the peptide environments surrounding Site 0 (left) and Sites 1 and 2 of View 2. The local color and radius of the cartoon representation reflect the average experimental thermal parameters (B factor) of each amino acid.

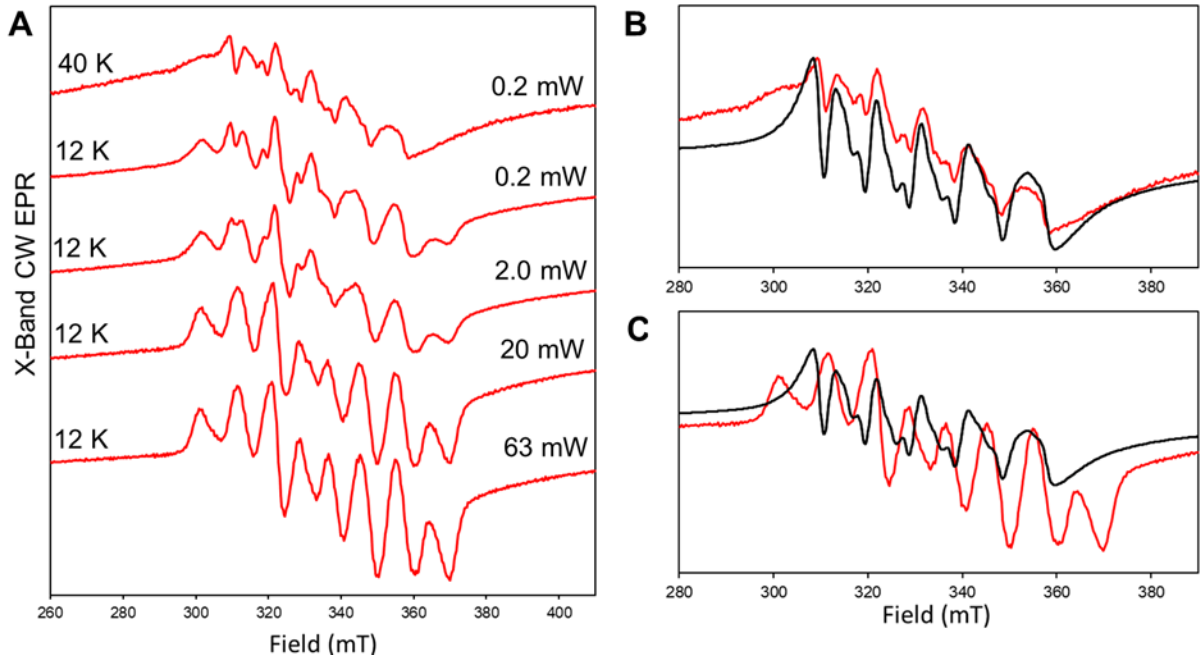


**Figure S5.** (**A**) Normalized X-band CW EPR spectra of $^{Mn}$AibH1H2 in 20 mM CHES pH 8.9 (red) collected at the listed powers and temperatures. Overlays of the X-band CW EPR spectrum of an aqueous 250 µM $MnCl_2$ standard (black, 5K, 2 mW) and **(B)** $^{Mn}$AibH1H2 at 40 K and 0.2 mW (red) or **(C)** $^{Mn}$AibH1H2 at 12 K and 63 mW (red).

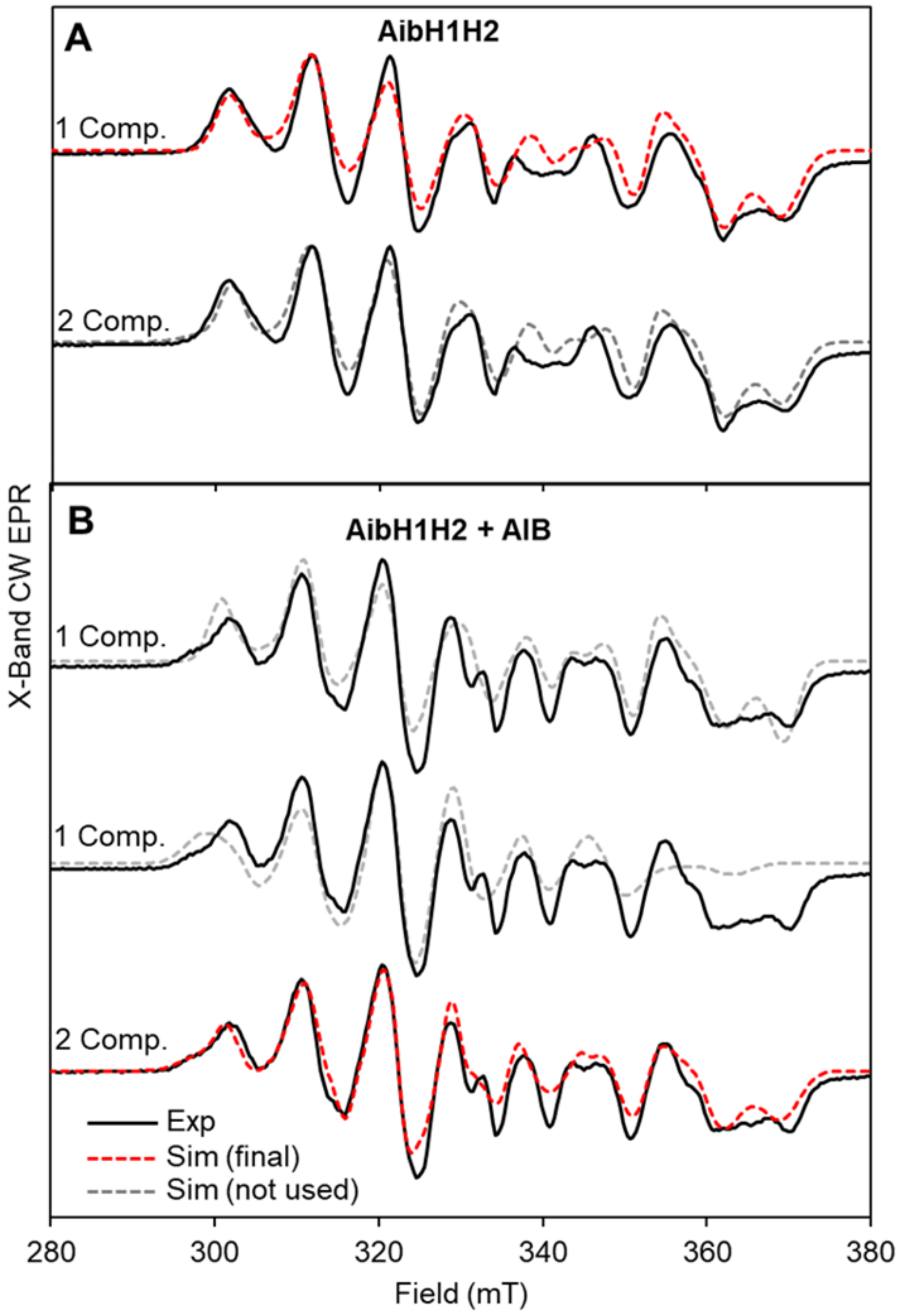


**Figure S6.** Different simulations of X-band EPR spectra of $^{Mn}$AibH1H2 in 20 mM CHES pH 8.9 without (**A**) and with (**B**) 100 mM AIB. Experimental spectra are shown as solid black lines, the best simulations chosen for the main text are in red dashed lines, and attempted simulations that were not used are in gray dashed lines. For the sample without AIB (**A**), the fit is similar with a single- or multiple-component simulation, and therefore a single-component simulation was chosen. For the sample with AIB (**B**), simulations with only one component failed to simultaneously model the new feature around 297 mT and the high-field features.

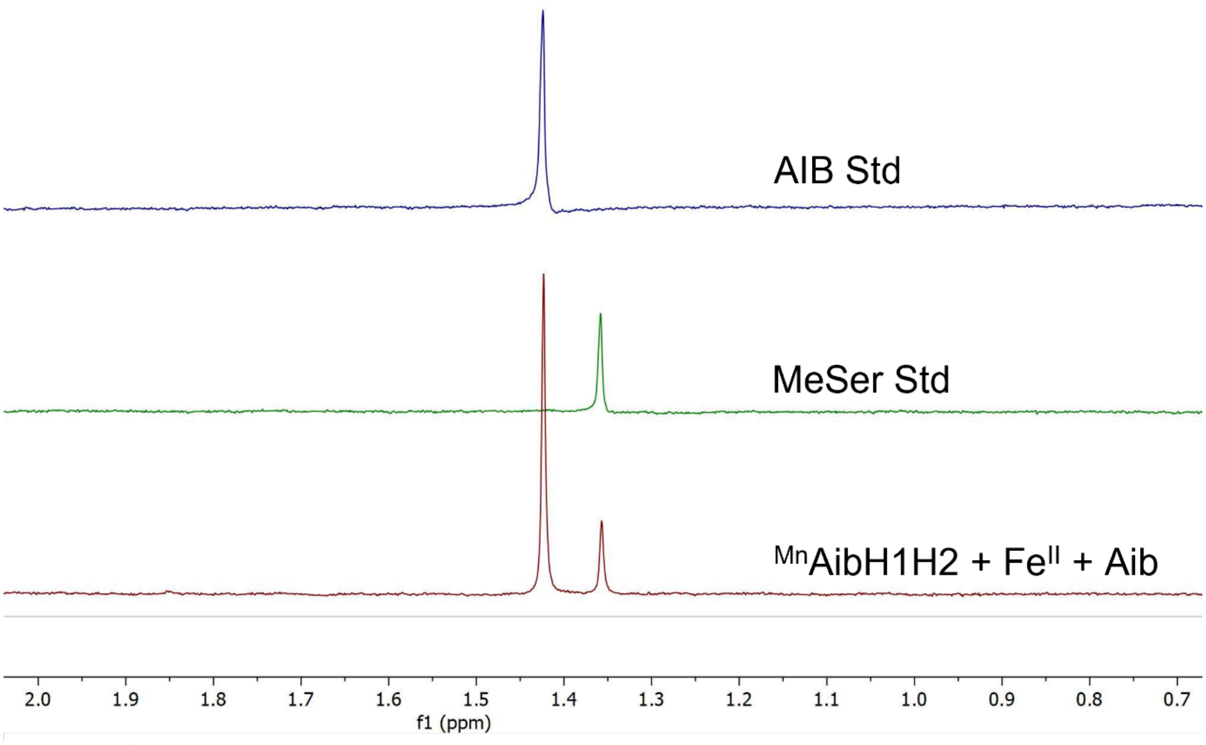


**Figure S7.** $^{1}$H NMR ($D_2O$, 400 MHz, 22 ºC) spectroscopic characterization of enzymatic reaction products. Enzymatic reactions were performed as described above using 500 µL total reaction volume and employing 200 µM $^{Mn}$AibH1H2, 200 µM $Fe^{II}$ metal, 5 mM ascorbate and 5 mM AIB. Following 3 h of aerobic agitation, the protein was removed by centrifugation in a 30 kDa Amicon® centrifugal unit. A 400 µL aliquot of the resultant flowthrough was added to 100 µL of $D_2O$ for $^{1}$H NMR spectroscopic analysis.

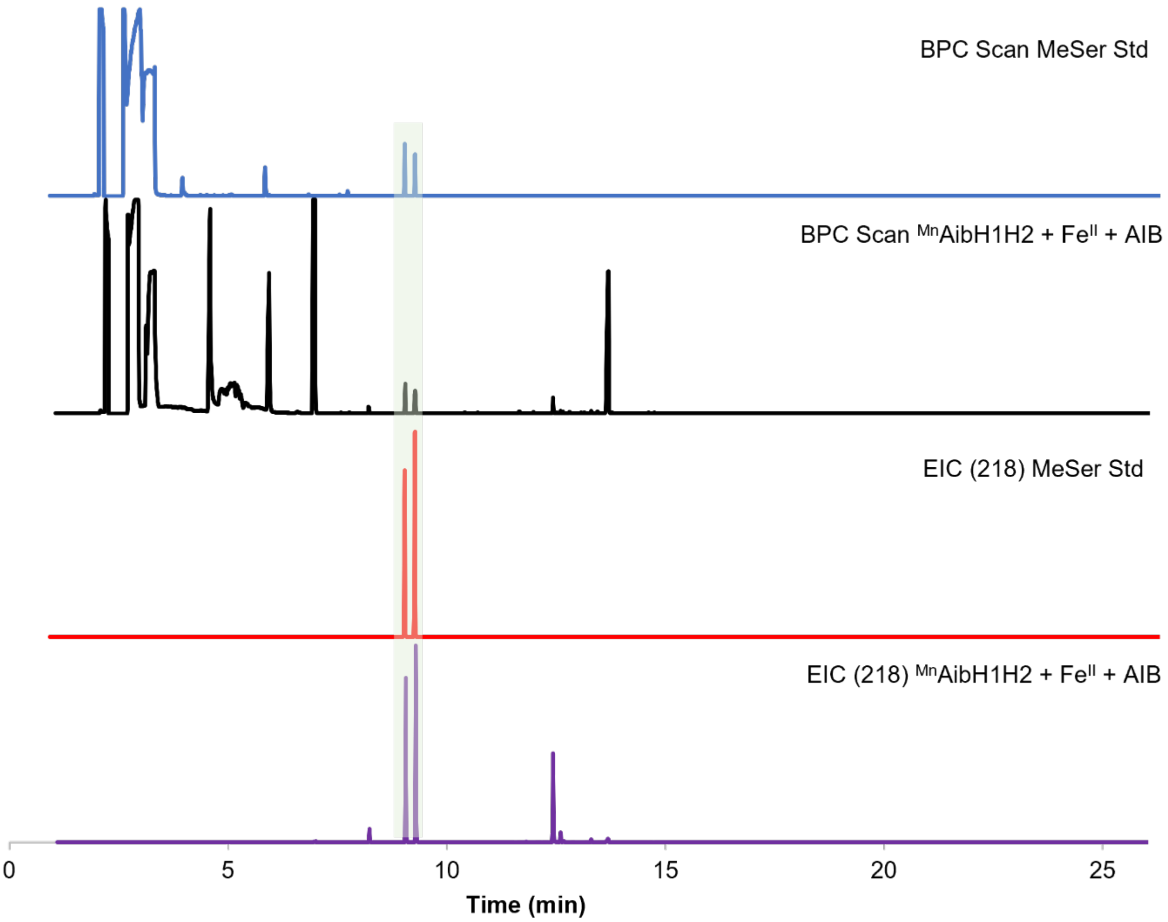


**Figure S8.** Complete base peak chromatogram (BPC) and the complete extracted ion chromatogram (EIC) filtered at m/z = 218 for a representative enzymatic assay of $^{Mn}$AibH1H2 and 1 equiv $Fe^{II}$ and a MeSer standard. The peaks for MeSer (elution time 9.3 minutes) and the internal standard serine (9.1 minutes) are highlighted in green.

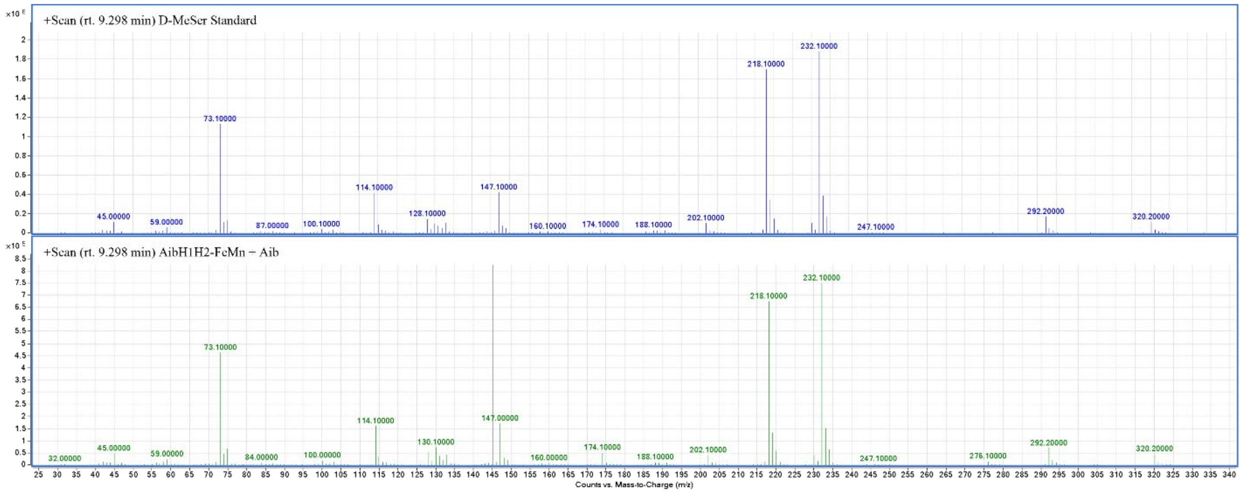


**Figure S9.** Mass spectra at 9.3 min retention time (Fig S8) for (top) the *D*-MeSer standard and (bottom) the product of the enzymatic reaction of $^{Mn}$AibH1H2 and 1 equiv $Fe^{II}$.

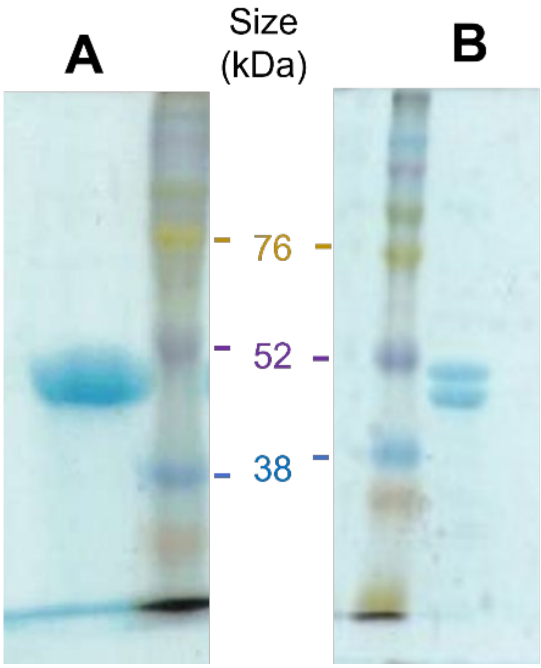


**Figure S10.** Representative SDS-PAGE gels of (A) full-length AibH1H2 and (B) AibH1H2 with a Δ17-24 truncation on AibH1.

| | Site 0 | Site 1 | Site 2 | $^{35}$Cys(S) AibH1 | $^{116}$Cys(S) AibH1 | $^{107}$Cys(S) AibH2 | $^{131}$Cys(S) AibH2 | $^{328}$Cys(S) AibH2 |
|---|---|---|---|---|---|---|---|---|
| ***Chain A*** | | | | | | | | |
| Mn Edge | **18.8** | - | - | 4.5 | 3.4 | - | - | - |
| Fe Difference | 2.2 | - | - | 1.6 | 2.3 | - | - | - |
| ***Chain B*** | | | | | | | | |
| Mn Edge | - | 5.3 | **8.1** | - | - | 2.2 | 2.8 | 2.6 |
| Fe Difference | - | **7.4** | 2.6 | - | - | 2.1 | 2.9 | 3 |
| ***Chain C*** | | | | | | | | |
| Mn Edge | **17.9** | - | - | 2.2 | 3.1 | - | - | - |
| Fe Difference | <2 | - | - | 2.1 | 2.4 | - | - | - |
| ***Chain D*** | | | | | | | | |
| Mn Edge | - | **12.4** | **7** | - | - | 2.7 | 3.7 | 4 |
| Fe Difference | - | 3.5 | <2 | - | - | 2.3 | 2.1 | 3.2 |

**Table S1.** Anomalous difference peak heights for structure 8FUN reported in σ values. Boldened values reflect our atomic assignments. Peak heights at the five cysteine sulfur atoms present in AibH1H2 are included to gauge the intrinsic noise level in this dataset.

**Table S2:** Simulation parameters for the EPR spectra discussed in the main text.

| | | | | | | | $Mn^{III}$ | | | | |
|---|---|---|---|---|---|---|---|---|---|---|---|
| | | $g_1$ | $g_2$ | $g_3$ | $g_{iso}$[a] | $g_{aniso}$[b] | $A_1$ | $A_2$ | $A_3$ | $A_{iso}$ | $A_{aniso}$ |
| $^{NA}Fe$ (1 M Tris) | | 2.033 | 2.029 | 2.024 | 2.029 | 0.007 | 303 | 245 | 375 | 308 | 101 |
| $^{57}Fe$ (1 M Tris) | | 2.033 | 2.029 | 2.024 | 2.029 | 0.007 | 303 | 245 | 375 | 308 | 101 |
| No Aib (20 mM CHES) | | 2.042 | 1.984 | 1.928 | 1.985 | 0.086 | 298 | 259 | 242 | 266 | 48 |
| 100 mM Aib (20 mM CHES) | Sys1[c] | 2.042 | 1.984 | 1.928 | 1.985 | 0.086 | 298 | 259 | 242 | 266 | 48 |
| | Sys2 | 2.055 | 2.044 | 2.020 | 2.039 | 0.030 | 264 | 199 | 385 | 283 | 154 |

| | $Fe^{III}$ | | | | |
|---|---|---|---|---|---|
| | $A_1$ | $A_2$ | $A_3$ | $A_{iso}$ | $A_{aniso}$ |
| $^{57}Fe$ (1 M Tris) | -65 | -66 | -68 | -66 | 2.5 |

[a]The isotropic g- and A- terms are an average of the individual values. [b]The anisotropy in the g- and A- tensors is given as the difference between the unique component and the average of the other two components. [c]The ratio of Sys1:Sys2 is 3.63:0.51

**AibH1**

ATGC**CCATGG**GC**CATCATCATCATCATCAC**AGCGGC**GAAAACTTATACTTTCAATCCGGT**GGCATGGTTGCGC
CCACCTCGAATCCAGGGGTACCGGATGAATTGGACGGTGTACCTGCGGTCGTGGACTGCGACGTACACGCGGT
TTTGCCGTCCCCTCATTCTCTGATTCCATATTTGGACGAGTACTGGGCCGACCAGCTTGTCGCACAGTTAGCTCC
TACGTATGAACCTAACTACCATCCGCGCGGATCCGCAATAGCACAACACTCAGACGCGAGTGTTGATGAGAACG
GCCGTGCGGCCACCACGGCTGAGAACTTGGTGAAGGACGTATTCGCCGACGGCTTTACGGACTTTGCTGTGGT
CAACTGCCTTTACGGCGTGCAACAAATACACCAACCTCGGCGTGAGATGGCACATGCTCGGGCTTTGAACCACT
GGATAGCGAATGAGTGGCTGGATAAAGACGATCGTCTGAGAGCCAGTATCGTGGTTCCCCAAGGGAGCCCTCG
GGCTGCCGCTGAGGAGATCGATTTTTGGTCTGGAGACAAGAGATTCGTACAAGTATTGCTGTTAGGGCAGAGCG
AACTGCTTTACGGAAGAGAAATCAATTGGCCCATCTGGGAAGCGGCAGAAGCTGCCGGGCTTCCCGTAACTTTA
CATATTGGTGGCGTTTTTCGTCAGGCTCCAACTAGCGTGGGATGGCCCGCATCACATTTGGAATGGTACGTGGG
ACAGCAGAGTAATATCGAGGCCCAGCTGAACTCGATCATTAGCGAAGGGATTCTTCAAAAATTTCCAAAAACAAA
GATCTTATTAAGTGAACTTGGCTTCAACTGGCTTCCACCGTTTATGTGGAAATTTGACAAATTATGGAAAAGCTAT
AGACCCGATATTCCTTGGGTTCAGGAATCGCCTTTGGAATTGATACGCGAGCACGTTCGGGTTACCACCAGTCC
AAGCGATGGCGCAGAGGAGGCAGGCCGCCTTGACTCTATCGTGGACCGGTTAGGAAGTGACCGCATGTTAGTT
TACAGTTCTGATTATCCCCACAAGCACCACTCAGGTCCAAGAGACATTGAAAATGGGACCCACAGCCCTGAATTA
TTGGACCGGATATACCGTCGGAACGCCTTCGATCTTTACAACCTTGTAGTGCCCTCCCCCGGAAAAGTGGGTTA
A**AAGCTT**ATGC

**AibH2**

**GGAATTCCATATG**ACGATAATCGAGCATGGGTCCCTGGGTACATTACCGGCGCCAAGTGTCACCACCGGGATC
GTCGACGCGGATATCCATCCAGTACCACAGGACGGCGCGCTTGAGCCATATCTGGATGATCGTTGGAAAAAACA
CATCAGAGAATACGGAGTCCGGACGACCACAGGCCTGCAGTTTATTTCAGAGTACCCACAAATGTATGGTGGGG
CAATGAGAGCAGACGCCTGGCCTGAATCGGGGTATCCTGGGAGCGACCGTGAGCTTCTGCGGACACAATTGTT
GGACAAACACAATATACAACTGGGGGTGTTGCAGTGCCTTGCCCCCGGCGGGCAAACCCTGAATCCGGCTGGA
CAGGCCTTAAACCAAGAACTTGCCGCTGCCCTTTGTCGTGCTACTAATGACTGGCAATTGGAGCACCTTGTGTAC
CCCGACCCACGGATGAGAGCTGCAATCCCTGTGACTTTTGAGACCCCTGACTACGCAGTCGCTGAAATAGAACG
TGTGGGAGCAGATCCAGGTGTCGTTGCCGTTTTAGGGACGAGCAAAACGTTGGAGCCTCTGGGAAGTCGCAAG
TACTGGCCGATCTACGAGGCGTCCGTCGCGCAGAATCTTCCGATACAGTTCCACTTGTCGCAAGGCGGGGGAC
ACGCTAATACAGGAACGGGATGGACCTCATATCATACGGAATATCACACAGGACATGTTCAATCTTTTCAATCGC
AGTTACTGAGTCTTGTATTATCAGGCACTTTCGACCGCTTTCCAACCCTTAAAGTTATGTTCGTGGAGGGTAATGT
GGCTCATTTCGCTCCACTTATCCAACGCATGGATTATACTTGGGAGACGCTTCGCGGAGAGCTGCCAGACCTGC
AGCGTAAGCCATCTGAGTACATACGTGATCACATTTGGGCGAGCACCCAGCCCATAGATGAGCCTGAAAAGCCG
GAGCACTTAGCGGAGTTATTAGAAGAGTTCTGCGGGGACAATGTCGTTTTCGCAACAGATTACCCCCACTTTGAT
TTCGATGACCCTGAGACTGCGTTTCCACGCTCGTTTCCTGTCGACCTTAGAGATAAGATCTTACGCGGAAATGGT
ATGCGCTTTTTTGGCGTAACGAACCAGGCTGATTAA**CTCGAG**ATGCG

**Table S3.** Gene blocks (5’ to 3’) used for AibH1H2 plasmid construction. Restriction enzyme recognition sequences are colored red and underlined, linker bases and the stop codon (TAA) are colored red, the 6x-His tag is underlined and bolded, and the TEV protease recognition sequence is boldened.

| Name | Sequence (5'→3') | Application |
|---|---|---|
| LacI-F | GGCATACTCTGCGACATCGT | Sequencing |
| pAC-Md-F | GTCGAACAGAAAGTAATCGTATTG | Sequencing |
| T7 term | GCTAGTTATTGCTCAGCGG | Sequencing |
| pAC-T7-1R | CGTATTAATTTCGATTATGCGG | Sequencing |
| T7 | TAATACGACTCACTATAGGG | Sequencing |
| DuetUP | TTGTACACGGCCGCATAATC | Sequencing |
| DuetDOWN | GATTATGCGGCCGTGTACAA | Sequencing |
| AibH1_Δ17-24-F | CTTATACTTTCAATCGAATCCAGGGGTACCGGATGAATTGG | Mutagenesis |
| AibH1_Δ17-24-R | CTGGATTCGATTGAAAGTATAAGTTTTCGCCGCTGTGATG | Mutagenesis |

**Table S4.** Primers used in this study.

| Name | Molecular Weight (kDa) AibH1 | AibH2 | AibH1H2 | $\varepsilon_{280}$ ($M^{-1}$ $cm^{-1}$) | Application |
|---|---|---|---|---|---|
| pET-Duet-AibH1H2 | 43.9 | 42.2 | 86.1 | 72365 | EPR, catalytic activity assays, crystal structure of $^{LB}$AibH1H2 (PDB: 8FUL) |
| pACYC-AibH1H2 | 43.9 | 42.2 | 86.1 | 72365 | $^{Fe}$AibH1H2 crystals (PDB: 8FUM and 8FUO) |
| pET-Duet-AibH1H2-Δ17-24 | 41.3[a] | 42.2 | 83.5 | 71620 | $^{Mn*}$AibH1H2 + Fe crystal (PDB: 8FUN) |

[a]Size after removal of the $His_6$-tag by TEV cleavage

**Table S5.** Summary of plasmids used in this study.

| Crystal | Plasmid | Beamline | Energy (eV) | Metal[a] | Precipitant |
|---|---|---|---|---|---|
| 8FUN | pET-Duet-AibH1H2-Δ17-24 | ALS 8.2.2 | 6550, 7132, 7092 | $(NH_4)_2Fe(SO_4)_2(H_2O)_6$ | 0.16 M $MgCl_2$, 0.08 M Tis-HCl pH 8.7, 24% PEG4000, 50 mM AIB, 20% glycerol |
| 8FUM | pACYC-AibH1H2 | SSRL 9-2 | 12658 | $MnCl_2{\cdot}4H_2O$ | 0.16 M $MgCl_2$, 0.08 M Tis-HCl pH 8.7, 20% PEG4000, 50 mM AIB |
| 8FUO | pACYC-AibH1H2 | SSRL 9-2 | 7132 | *none* | 0.16 M $MgCl_2$, 0.08 M Tis-HCl pH 8.7, 20% PEG4000 |
| 8FUL | pET-Duet-AibH1H2 | ALS 5.0.2 | 12658 | *none* | 0.16 M $MgCl_2$, 0.08 M HEPES pH 8.5, 20% PEG4000, 20% glycerol, 10 mM DT |

[a]2 equivalents of metal added relative to [AibH1H2]

**Table S6.** Summary of crystallization conditions in this study.

| | $^{Fe}$AibH1H2 | $^{LB}$AibH1H2 | $^{Fe}$AibH1H2 | $^{Mn*}$AibH1H2 + 2 equiv $Fe^{II}$ | $^{Mn*}$AibH1H2 + 2 equiv $Fe^{II}$ | $^{Mn*}$AibH1H2 + 2 equiv $Fe^{II}$ |
|---|---|---|---|---|---|---|
| **PDB ID** | 8FUO | 8FUL | 8FUM | 8FUN | | |
| **Data Collection** | | | | | | |
| Wavelength | 1.738 Å | 0.979 Å | 0.979 Å | 1.738 Å | 1.748 Å | 1.893 Å |
| Resolution range (Å) | 79.74 - 2.43 (2.52 - 2.43) | 47.28 - 2.29 (2.33 - 2.29) | 45.58 - 1.48 (1.53 - 1.48) | 46.0 - 2.24 (2.29 - 2.24) | 46.0 - 2.24 (2.29 - 2.24) | 46.0 - 2.50 (2.58 - 2.50) |
| Space group | I 2 2 2 | I 1 2 1 | I 1 2 1 | I 2 2 2 | I 2 2 2 | I 2 2 2 |
| Unit cell | 84.78 150.57 234.72 90 90 90 | 82.77 231.40 145.00 90 92.17 90 | 82.59 232.64 147.64 90 92.77 90 | 84.14 148.92 234.09 90 90 90 | 84.19 148.83 233.87 90 90 90 | 84.22 148.83 233.89 90 90 90 |
| Total reflections | 650852 (44054) | 373023 (18655) | 1102416 (117933) | 800957 (32071) | 800785 (30431) | 615729 (47432) |
| Unique reflections | 55038 (5260) | 119390 (11703) | 431064 (42532) | 70480 (6547) | 70089 (3953) | 51127 (4361) |
| Multiplicity | 11.8 (10.3) | 3.1 (3.1) | 2.8 (2.6) | 11.4 (8.0) | 11.4 (7.7) | 12.0 (10.9) |
| Completeness (%) | 96.7 (93.9) | 97.9 (96.1) | 93.72 (92.58) | 99.3 (93.46) | 98.9 (87.5) | 99.8 (99.6) |
| Mean I/sigma(I) | 18.3 (3.0) | 8.1 (4.1) | 7.2 (1.2) | 5.2 (1.8) | 5.6 (1.8) | 7.7 (3.0) |
| Wilson B-factor | 49.9 | 46.15 | 12.7 | 25.25 | | |
| R-merge | 0.085 (0.738) | 0.08 (0.202) | 0.085 (0.818) | 0.486 (1.504) | 0.487 (1.566) | 0.267 (0.805) |
| R-meas | 0.089 (0.777) | 0.108 (0.281) | 0.105 (1.027) | 0.527 (0.809) | 0.534 (1.799) | 0.290 (0.886) |
| R-pim | 0.026 (0.236) | 0.061 (0.154) | 0.060 (0.608) | 0.154 (0.585) | 0.153 (0.626) | 0.082 (0.263) |
| CC1/2 | 0.999 (0.810) | 0.989 (0.915) | 0.996 (0.355) | 0.901 (0.293) | 0.916 (0.337) | 0.972 (0.753) |
| Anom Completeness | 97.3 (93.4) | n/a | n/a | 98.6 (84.8) | 98.4 (83.2) | 99.7 (99.2) |
| Anom Multiplicity | 6.1 (5.3) | n/s | n/a | 5.8 (4.2) | 5.8 (4.0) | 6.2 (5.5) |
| **Refinement** | | | | | | |
| Reflections used in refinement | 55032 (5259) | 119391 (11704) | 430996 (42524) | 70481 (6548) | | |
| Reflections used for R-free | 2000 (191) | 2000 (191) | 1414 (140) | 2000 (186) | | |
| R-work | 0.188 (0.264) | 0.1894 (0.3813) | 0.159 (0.253) | 0.1774 (0.2656) | | |
| R-free | 0.250 (0.347) | 0.2563 (0.4716) | 0.184 (0.250) | 0.1979 (0.2837) | | |
| Number of non-hydrogen atoms | 11658 | 23276 | 25133 | 12070 | | |
| macromolecules | 11347 | 22658 | 22812 | 11433 | | |
| ligands | 14 | 25 | 180 | 16 | | |
| solvent | 297 | 593 | 2141 | 621 | | |
| Protein residues | 1430 | 2852 | 2859 | 1436 | | |
| RMS(bonds) | 0.009 | 0.009 | 0.009 | 0.012 | | |
| RMS(angles) | 1.3 | 1.32 | 1.36 | 1.4 | | |
| Ramachandran favored (%) | 94.51 | 94.98 | 96.96 | 95.94 | | |
| Ramachandran allowed (%) | 5.07 | 4.84 | 2.79 | 3.57 | | |
| Ramachandran outliers (%) | 0.42 | 0.18 | 0.25 | 0.49 | | |
| Rotamer outliers (%) | 3.49 | 2.33 | 1.03 | 1.48 | | |
| Clashscore | 5.22 | 5.69 | 3.24 | 4.01 | | |
| Average B-factor | 52.91 | 48.49 | 17.46 | 27.6 | | |
| macromolecules | 53.21 | 48.6 | 16.9 | 27.59 | | |
| ligands | 58.9 | 62.06 | 35.09 | 38.64 | | |
| solvent | 41.15 | 43.37 | 21.87 | 27.64 | | |

**Table S7:** X-ray data collection and refinement statistics. Numbers in parenthesis correspond to statistics of the highest resolution shell.
